# Optical measurement of physiological sodium currents in the axon initial segment

**DOI:** 10.1101/2020.07.20.211839

**Authors:** Luiza Filipis, Marco Canepari

**Author notes:** Author contributions: L.F. conducted the experiments, performed the analysis and the modeling and wrote the paper; M.C. designed the method, the experiments and the modelling and wrote the paper.

## Abstract

In most neurons of the mammalian central nervous system, the action potential (AP) is triggered in the axon initial segment (AIS) by a fast Na^+^ current mediated by voltage-gated Na^+^ channels. The intracellular Na^+^ increase associated with the AP has been measured using fluorescent Na^+^ indicators, but with insufficient resolution to resolve the Na^+^ current in the AIS. In this article, we report the critical improvement in temporal resolution of the Na^+^ imaging technique allowing the direct experimental measurement of Na^+^ currents in the AIS. We determined the AIS Na^+^ current, from fluorescence measurements at temporal resolution of 100 *µ*s and pixel resolution of half a micron, in pyramidal neurons of the layer-5 of the somatosensory cortex from brain slices of the mouse. We identified a subthreshold current before the AP, a fast inactivating current peaking during the rise of the AP and a persistent current during the AP repolarisation. We correlated the kinetics of the current at different distances from the soma with the kinetics of the somatic AP. We quantitatively compared the experimentally measured Na^+^ current with the current obtained by computer simulation of published NEURON models and we show how the present approach can lead to the correct estimate of the native behaviour of Na^+^ channels. Thus, it is expected that the present method will be adopted to investigate the function of different channel types under physiological or pathological conditions.

## INTRODUCTION

In most neurons of the mammalian central nervous system the action potential (AP) is initiated in the proximal unmyelinated part of the axon (Bean, 2007), the so-called axon initial segment (AIS). Thus, the shape of the AP and the firing properties that distinctively characterise different neuronal types are determined by the synergistic behaviour of voltage-gated channels and associated proteins expressed in this fundamental compartment of polarised neurons (Rasband, 2010). Among these channels, voltage-gated Na^+^ channels (VGNCs) are the AP trigger since they mediate the transient inward current underlying the rapid rising phase of the AP (Kole & Stuart, 2012). Gaining the ability of precisely recording Na^+^ currents at the AIS is therefore crucial in the investigation of the AP generation under physiological conditions or in the case of the frequent neurological disorders caused by AIS dysfunction (Wimmer *et al.* 2010). This goal, however, has been thus far beyond available experimental techniques.

Fast Na^+^ currents can be recorded in brain slices using standard electrode techniques (Yue *et al.* 2005; Astman *et al.* 2006). These measurements, that do not contain any information on the spatial profile of the current, are achieved by “clamping” the membrane potential (V_m_) and therefore can be only indirectly correlated with the physiological AP. In contrast, axonal Na^+^ influx associated with real APs can be measured at the site of origin by imaging the fluorescence of a Na^+^ indicator (Kole *et al.* 2008; Fleidervish *et al.* 2010; Baranauskas *et al.* 2013). Yet, the most important studies using this approach were performed using sodium-binding benzofuran isophthalate (SBFI, Minta & Tsien, 1989) as indicator, at acquisition rates insufficient to precisely reconstruct the kinetics of the Na^+^ influx during the AP. While faster Na^+^ fluorescence measurements are limited by the sensitivity of this indicator, a significant improvement in the signal-to-noise ratio (SNR) can be obtained by using the indicator Asante NaTRIUM Green-2 (ANG-2, Miyazaki & Ross, 2015; Miyazaki *et al.* 2019) or the commercially available Ion NaTRIUM Green-2 (ING-2, Ion Indicators). In particular, with ∼5% change of fluorescence corresponding to 1 mM (Lamy & Chatton, 2011), these indicators allow resolving axonal Na^+^ transients associated with a single AP with a resolution of a few microns (Miyazaki & Ross, 2015). With these premises, using recent commercially available technology and a system developed in our laboratory, we measured Na^+^ fluorescence with a temporal resolution of 100 *µ*s to correlate the kinetics of Na^+^ influx with that of the AP. We used the layer-5 (L5) pyramidal neuron of the somatosensory cortex of the mouse to develop this novel technique. From these fast Na^+^ concentration measurements, we reconstructed the Na^+^ current associated with the AP by calculating the time derivative of the fluorescence transient, similarly to what was done to reconstruct fast Ca^2+^ currents (Jaafari *et al.* 2014; Jaafari *et al.* 2015; Jaafari & Canepari, 2016). Notably, in a fraction of cells, a subthreshold component of the Na^+^ current preceding the AP was detected above the photon noise, while the current component associated with the AP was always composed by a fast and a slow current. We could therefore reconstruct the quantitative profile of the Na^+^ current along the ∼40 *µ*m long AIS of the L5 pyramidal neuron of the mouse and compare these currents with those predicted by NEURON simulations.

## METHODS

### Slice preparation and solutions

Experiments were ethically carried out in accordance with European Directives 2010/63/UE on the care, welfare and treatment of animals. Procedures were reviewed by the ethics committee affiliated to the animal facility of the university (D3842110001). We used 21-35 postnatal days old mice (C57BL/6j) purchased from Janvier Labs (Le Genest-Saint-Isle, France). Animals were anesthetised by isofluorane inhalation and the entire brain was removed after decapitation. Brain slices (350 *µ*m thick) were prepared using the following procedure. After separating the entire brain from the skull, the cerebellum was dissected away and discarded. The two hemispheres of the brain were separated with a blade and glued from the sagittal plane on a custom made tissue holder with an angle of 15 degrees from the coronal plane. Slices were cut with the blade using a Leica VT1200 vibratome (Wetzlar, Germany). Following the same protocols used in the laboratory for other preparations (Jaafari *et al.* 2014; Jaafari & Canepari, 2016; Ait Ouares *et al.* 2019; Ait Ouares & Canepari, 2020), slices were incubated at 37°C for 45 minutes and maintained at room temperature before use. Slices with L5 pyramidal neurons in the somato-sensory cortex having an axon parallel to the slice surface, as visualised with transmitted light, were selected. The extracellular solution contained (in mM): 125 NaCl, 26 NaHCO_3_, 1 MgSO_4_, 3 KCl, 1 NaH_2_PO_4_, 2 CaCl_2_ and 20 glucose, bubbled with 95% O_2_ and 5% CO_2_. The intracellular solution contained (in mM): 125 KMeSO_4_, 5 KCl, 8 MgSO_4_, 5 Na_2_-ATP, 0.3 Tris-GTP, 12 Tris-Phosphocreatine, 20 HEPES, adjusted to pH 7.35 with KOH. In V_m_ imaging experiments, cell membranes were loaded with the voltage-sensitive dye D-2- ANEPEQ (JPW1114, 0.2 mg/mL, Invitrogen, Carlsbad, CA) for 30 minutes using a first patch clamp recording and then re-patched a second time with dye free solution as previously described (Canepari *et al.* 2008). In all other experiments, the Na^+^ indicator ING-2 (IonBiosciences, San Marcos, TX) was added to the intracellular solution at the concentration of 0.5 mM and recordings were started 20 minutes after achieving the whole cell configuration. While V_m_ imaging can be combined with Ca^2+^ imaging (Vogt *et al.* 2011a) or photorelease (Vogt *et al.* 2011b), The overlap in the emission spectra of the two indicators did not allow the combination of V_m_ and Na^+^ imaging the same cells.

### Electrophysiology and pharmacology

Patch-clamp recordings were made at the temperature of 32-34°C using a Multiclamp 700A (Molecular Devices, Sannyvale, CA) and electrical signals were acquired at 20 kHz using a USB-6221 board (National Instruments, Austin, TX) and a custom-written code written in MATLAB. The measured V_m_ was corrected for junction potential (−11 mV) as previously estimated (Canepari *et al.* 2010). APs were elicited in current clamp mode by 5 ms current pulses of 1.5-2.5 nA through the patch pipette. The bridge was also corrected offline by using the recorded injected current. Tetrodotoxin (TTX, purchased from Tocris, Bristol, UK) and huwentoxin-IV (HwTx-IV, purchased from Smartox Biotechnology, Saint Egrève, France) were dissolved in extracellular solution and delivered through a pipette positioned near the AIS by applying a gentle positive pressure. The timing of illumination, the start of the camera acquisition and the timing of the somatic current injection were precisely controlled by a Master-9 pulse stimulator (A.M.P.I., Jerusalem, Israel). The alignment of the somatic recording with the camera frames was done using an OptoFlash LED (Cairn Research, Faversham, UK) since this device has an illumination rise of ∼1 *µ*s.

### Optical recordings and initial analysis

Optical recordings were done using a modified version of a previously used system (Filipis *et al.* 2018). The imaging set-up consisted on an upright Scientifica SliceScope microscope equipped with a motorised XY translation stage and PatchStar manipulators (Scientifica, Uckfield, UK), and a 60X Olympus water immersion objective (NA = 1). Fluorescence was excited using a 520 mW line of a LaserBank (Cairn Research) band-pass filtered at 517 ± 10 nm and directed to the preparation using a 538 nm long-pass dichroic mirror. The size of the illumination spot was adjusted using a custom-made telescope. ING-2 Fluorescence emission was band-pass filtered at 559 ± 17 nm, demagnified by 0.5X and acquired with a DaVinci 2K CMOS camera (SciMeasure, Decatur, GA) at 10 kHz with a pixel resolution of 30×128. 80 frames (corresponding to 8 ms) were acquired except in the recording associated with two consecutive APs where the number of frames was 110 (corresponding to 11 ms). In V_m_ imaging recordings, JPW1114 fluorescence was long-pass filtered at >610 nm before being acquired also at 10 kHz for 8 ms. Data were acquired using the software Turbo-SM (RedshirtImaging, Decatur, GA). For the extraction of the Na^+^ current, fluorescence from 3-6 trials including APs signals was averaged after carefully checking that APs were precisely superimposing.

### Calibration and compensation for longitudinal diffusion

ING-2 ΔF/F_0_ transients, spatially filtered and corrected for photo-bleaching, were converted in terms of Δ[Na^+^] in the following manner. Using an internal solution without Na_2_-ATP, ING-2 fluorescence at different NaCl concentrations (0, 2.5, 5, 7.5, 10, 12.5, 15 and 25 mM) was measured with the same apparatus used in the physiological recordings. We linearly fitted the fluorescence against the Na^+^ concentration and we established that ΔF/F_0_ = 1% corresponds to Δ[Na^+^] = 0.175 mM at initial [Na^+^] = 5 mM. In order to compensate for the Δ[Na^+^] contribution due to longitudinal diffusion, the following procedure was applied. From the general theory (Crank, 1975), the [Na^+^] at longitudinal position *x* and time *t* is given by the equation:

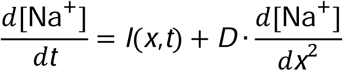

where *D* is the diffusion coefficient and *I*(*x,t*) is the Na^+^ influx at longitudinal position *x* and time *t*. Obviously, in the absence of diffusion (*D* = 0), the time derivative of [Na^+^] would correspond to *I*(*x,t*), but the diffusion coefficient for Na^+^ is D= 0.6 *µ*m^2^ ms^-1^ (Kushmerick *et al.* 1969). The compensating factor ([Na^+^]_diff_) to subtract from the experimental [Na^+^] is therefore given by the equation:

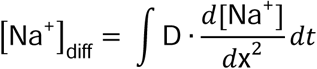

We estimated [Na^+^]_diff_ using the following procedure. First, the number of transient Na^+^ ions (*N*_*ions*_(*x,t*)) was obtained from the experimentally measured Δ[Na^+^] by estimating the volume of each compartment. Specifically, we applied a simple mask to the fluorescence image so that the pixels below a certain threshold were considered zero. Then, the diameter d(*x*) was obtained by using the number of consecutive positive pixels at the line *x* and an equation *d(x)* = *A·x*^*B*^ + *C* was fitted against the *x* axis to obtain a smooth realistic profile. It is important to point out that the direct measurement of the axon diameter is not possible with the present spatial resolution and its estimate, using the fit with a continuous realistic function, is therefore the best approximation to reduce the possible errors in the calculation of the Na^+^ current. Knowing the diameter of the axonal compartment at each plane at *x* position, the volume (*vol*) corresponding to each cylindrical compartment was calculated. The Δ[Na^+^] was first converted into the transient of ion density (Δ[N_ions_]) by multiplying by the Avogadro constant (N_AV_, 6.02·10^23^ mol^-1^) and *N*_*ions*_ was calculated as *vol*·Δ[N_ions_]. Then, the diffusion compensating factor was calculated using the gradient between *N*_*ions*_(*x,t-Δt*) and the concomitant *N*_*ions*_ in the two adjacent regions at the step/distance *Δx* from *x* (*N*_*ions*_(*x± Δx,t-Δt*)) using the equation:

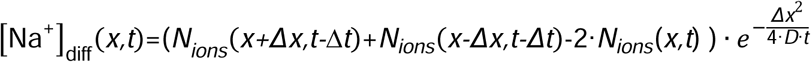

This expression neglects, as first approximation, the contributions of non adjacent (more far) regions that depend on the step/distance *Δx*. Since each value of [Na^+^] is essentially the average of 4 pixels (see the main text), we used *Δx* = 2 *µ*m. With this value, the contributions of non adjacent regions are negligible and the estimate of the compensating factor given above is realistic for the Na^+^ diffusion coefficient.

### Signals fit and calculation of the Na^+^ current

The compensated N_ions_ obtained with the above procedure was then converted into a charge surface density (δQ) by multiplying by the elementary charge constant (Q_e_, 1.6·10^−19^ C) divided by the surface of the disk at position *x*. The δQ signal was matched with the model function:

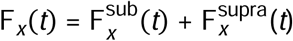

where the subthreshold and suprathreshold components were

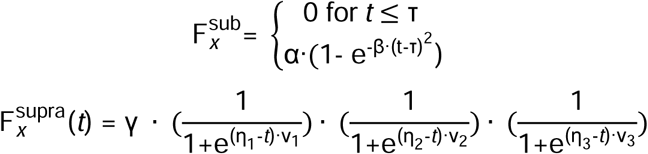

To maximise the likeliness between the model and the experimental trace the following procedure was applied. First, the parameters η_1_ and η_2_ were constrained to the exact time of the fast increase of the δQ transient and the parameter ν_1_ was arbitrarily set at 20 ms^-1^ corresponding to half sampling time of the fluorescence acquisition. Then, the experimental trace up to η_1_-2 was fitted with the subthreshold component using the Expectation-Maximization (EM) algorithm implemented in Matlab. In this way, the parameters α and β were constrained to the values obtained by this first fit. The entire experimental trace was fitted with the model function again using the Expectation-Maximization (EM) algorithm to determine the parameters γ, η_3_, ν_2_ and ν_3_. The I_Na_(*t*) signal at position *x* was eventually calculated as (F_*x*_(*t*)- F_*x*_(*t*-1))/*s*, where *s* is the sampling time (100 *µ*s). Since the estimated radius varied from 0.9 to 1.2 *µ*m at 5-10 *µ*m distance from the soma, and from 0.7 to 0.9 *µ*m at 30-35 *µ*m distance from the soma, the potential maximal error in the measurement of I_Na_ is ∼30% in either direction.

### Modifications of a published original NEURON model

The published model of Hallermann et al. (27), originally designed to calculate the metabolic cost of the AP in cortical pyramidal neurons, was modified to obtain AP waveforms and Na^+^ currents similar to the experimental ones. The original model can be found in the ModelDB database under the reference #144526 available at https://senselab.med.yale.edu/ModelDB/ShowModel?model=144526. VGNCs in the AIS were modified from the original model in the following manner. For “nax”, the activation parameter (Vshift) was shifted by 10 mV in the right direction and the inactivation parameter (Vshift_inact) was shifted by 25 mV also in the right direction. For “na channel”, the activation parameter (Vshift) was shifted by 17 mV in the right direction and the inactivation parameter (Vshift_inact) was shifted by 10 mV also in the right direction. The density of K^+^ channels was also modified as described later in the text.

### Statistical analysis

To quantify the spatial distribution of signals at three different distances from the soma (5-10 *µ*m, 15-25 *µ*m and 30-35 *µ*m), values obtained in the analysis were compared and significant differences in the spatial distribution were tested by performing the paired t-test with p <0.01 as discriminating threshold. Similarly, in pharmacological assessments, the block of the signals was tested by performing the paired t-test with p <0.01 as discriminating threshold to the signals in control condition and after addition of VGNC inhibitor. Finally, the correlation between two parameters was established by calculating the Pearson linear coefficient in Matlab and the related probability value for testing the null hypothesis with p <0.01 as discriminating threshold.

## RESULTS

### Optimisation of the spatial and temporal resolution of axonal Na^+^ imaging

To record axonal Na^+^ concentration changes in native neurons from brain slices, at the highest possible spatial and temporal resolution, we modified an imaging system originally developed for fast confocal Ca^2+^ imaging (Filipis *et al*. 2018) as illustrated in Fig. 1*A*. The light from a 1W - 520 nm laser, filtered at 517 ± 10 nm, was reflected by a 538 nm long-pass dichroic mirror to a 60X water-immersion objective, with numerical aperture equal to 1. The size of the illuminating spot was adjusted to either illuminate the whole field or an area of ∼30 *µ*m diameter using custom-made lenses. The emitted light was band-pass filtered at 559 ± 17 nm and images were demagnified by 0.5X before being acquired by a DaVinci2K CMOS camera. With this configuration, the field of view was covered by 512×512 pixels where the pixel size was ∼500 nm (Fig. 1*A*). The bottom images in Fig. 1*B* show a L5 pyramidal neuron, filled with 500 *µ*M of the Na^+^ indicator ING-2, illuminated either over the entire field of view or by the 30 *µ*m diameter spot only. We then placed the AIS in the recording position illuminated by the 30 *µ*m spot, covered by a sub-region of 30×128 pixels sampled at 10 kHz, as shown in the example of Fig. 1*C*. Fluorescence of each pixel was averaged on the *x-*axis, corresponding to a given distance from the soma calculated from the surface of the cell body, along the *y*-axis (Fig. 1*C*). We have previously estimated that for a fluorescent structure at 20-30 *µ*m from the slice surface, which is the case of an intact axon, fluorescence can be discriminated with a precision of 2-3 *µ*m (Filipis *et al.* 2018). Thus, to improve the signal-to-noise ratio (SNR) within the factual spatial resolution, we replaced the fluorescence F at each point ξ on the *x*-axe, with the Gaussian filtered fluorescence G(ξ) defined by the equation:

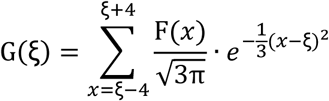

plotted at the top of Fig. 1*D*. Below this illustration, it is shown the fractional change of G(ξ) (ΔF/F_0_), associated with an AP and recorded at 10 kHz, calculated from the average of 4 trials. Photo-bleaching of the dye produced ∼10% decrease in fluorescence when illuminating a 30 *µ*m spot for 8 ms, as shown by the single recording without the AP (yellow trace in Fig. 1*D*). With the illumination confined to a small spot, fluorescence recovered over a minute by diffusion of the unbleached dye from non illuminated areas, permitting several sequential recordings with ∼1 minute from one to the next trial. To correct fluorescence from the effect of photo-bleaching, the recording without the AP was fitted by a tri-exponential function (dotted trace in Fig. 1*D*) which realistically models photo-bleaching of this specific indicator (Roder & Hille, 2014). This fit was subtracted to the ΔF/F_0_ signal to correct for the photo-bleaching effect (bottom red trace in Fig. 1*D*). This procedure reliably compensates the fluorescence decrease due to photo-bleaching since the mean ΔF/F_0_ of the first 8 samples, at all axonal positions, was consistently within one standard deviation from zero. With illumination confined to a small spot, fluorescence recovered over a minute by diffusion of the unbleached dye from non illuminated areas, permitting several sequential recordings with ∼1 minute from one to the next trial. Thus, the progressive photo-damage of the tissue was quantitatively assessed in order to establish how many trials can be performed before the physiological conditions are compromised. The analysis of the kinetics of the somatic AP, typically used to monitor the progress of photo-damage in V_m_ imaging (Canepari *et al.* 2008), is a reliable indicator of the photo-damage at the AIS as shown in the example of Fig. 2*A*, where 12 consecutive trials were performed. In this example, it is shown that the kinetics of the somatic AP and of the axonal Na^+^ transient were the same for the first and the fifth recording, whereas they were different for the twelfth recording. Thus, we investigated the change in the kinetics of the somatic AP, which mirrors that of the axonal Na^+^ signal, to assess how many trials at physiological conditions can be achieved. As shown in the analysis reported in Fig. 2*B*, the kinetics of the somatic AP is only slightly changing from that of the first AP recorded without illumination during the first 10 trials (with illumination), an effect that was consistently recorded in 6 cells tested in this way (Fig. 2*C*). We concluded that we can record the physiological Na^+^ influx in the AIS, associated with an AP, for at least 10 consecutive recordings, with a temporal resolution of 100 *µ*s and an effective spatial resolution of 2-3 *µ*m.

**Figure 1.**
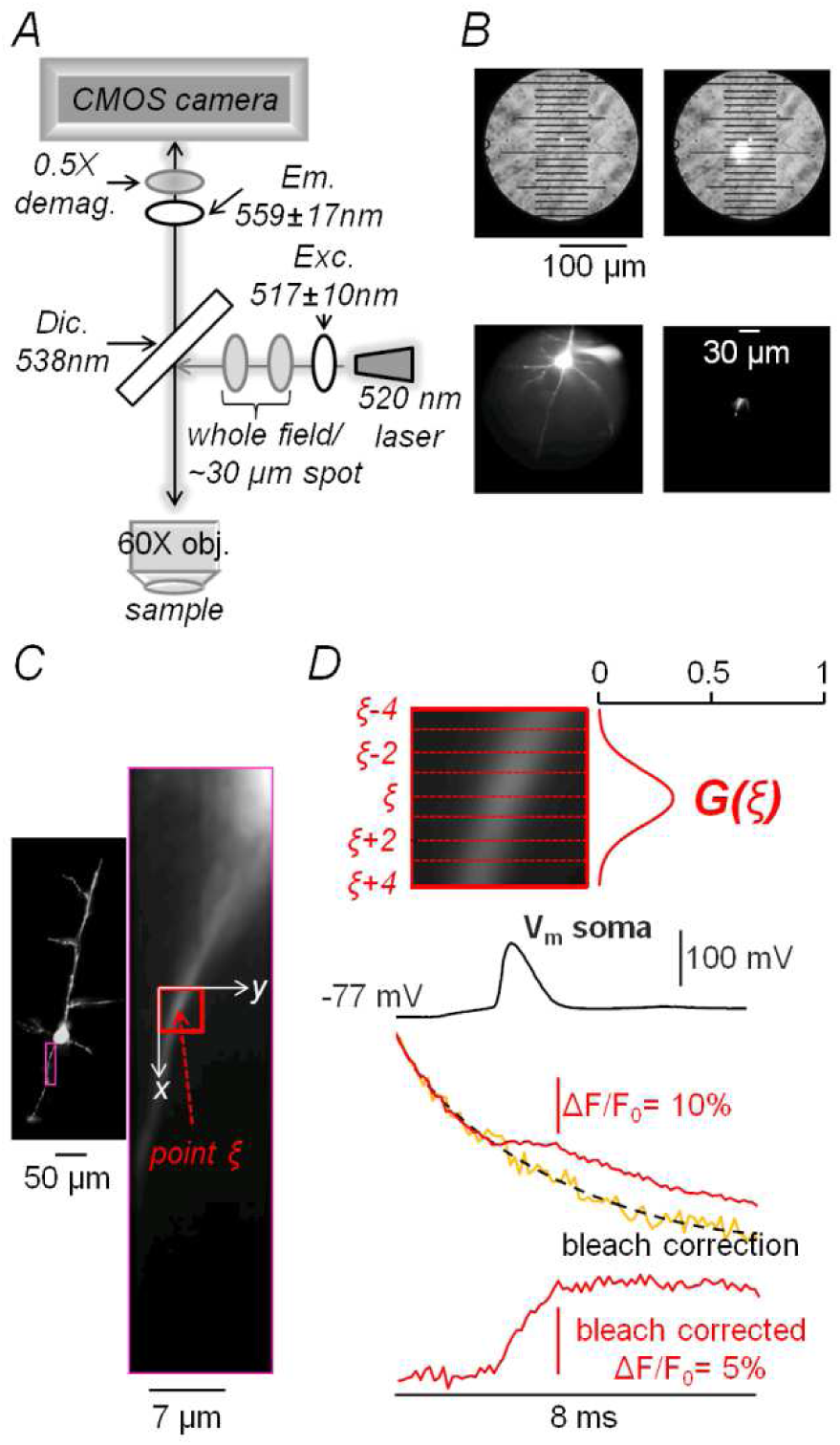
Experimental settings and principles of high-resolution Na^+^ imaging. *A*, design of the optical arrangement for high-resolution Na^+^ imaging. The 520 nm laser beam, band-pass filtered at 517 ± 10 nm, is either illuminating the whole field uniformly or a spot of ∼30 *µ*m using a 60X objective. The emitted light, passing through a 538 nm dichroic mirror, is band-pass filtered at 559 ± 17 nm. Images are demagnified by 0.5X to obtain a pixel resolution of ∼500 nm in the CMOS sensor. *B*, top, field of view corresponding to 512×512 pixels illuminated with transmitted light and focussing on a stage micrometer with 10 *µ*m steps graduation; the image on the right shows the laser illumination spot. Bottom, fluorescence from a L5 pyramidal neuron filled with 500 *µ*M ING-2 illuminated either uniformly (left) or with the laser spot (right). *C*, from another L5 pyramidal neuron, fluorescence in the recording position (30 × 128 pixels) including the laser spot; a region of interest is outlined by the red rectangle. *D*, the region of interest in panel *C* is zoomed and fluorescence from the 9 positions indicated with *ξ* is filtered with the plotted function *G(ξ)*. The black trace is an AP elicited by a 5 ms current pulse recorded in the soma. The red trace is the ING-2 ΔF/F_0_ from the region of interest recorded at 10 kHz and associated with the AP above. The yellow trace is a recording from the same region without stimulation. The dotted trace is the 3- exponential fit of the yellow trace performed to estimate the time-course of the bleach. On the bottom, the red trace is the ING-2 ΔF/F_0_ after bleach correction. Red traces are from averages of 4 trials.

**Figure 2.**
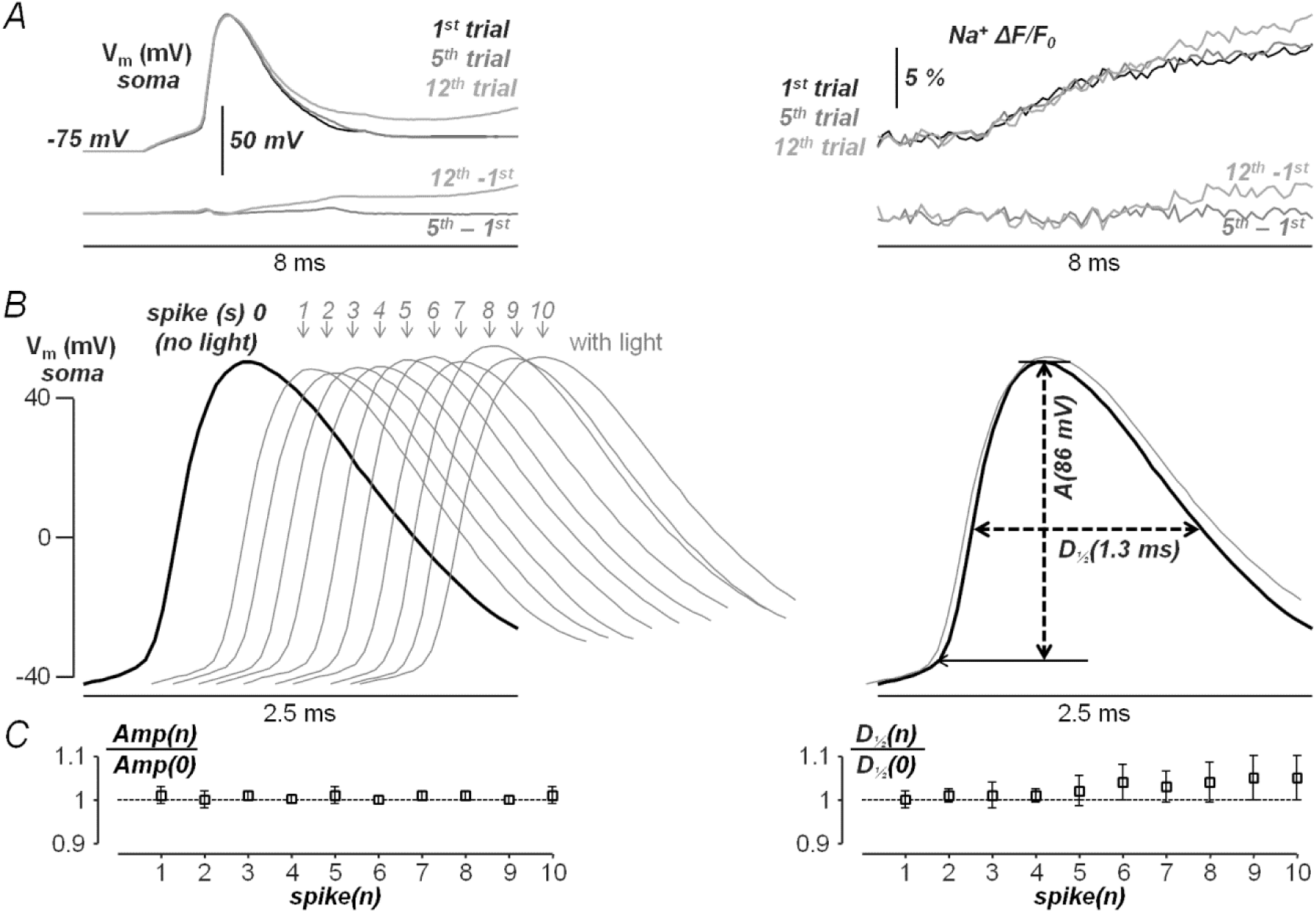
Amplitude and shape of the somatic action potential during 10 light exposures. *A*, left, somatic APs from sequential trials with light exposure of 8 ms and 1 minute interval between two consecutive recordings: black trace is the first trial; dark grey trace is the fifth trial; light grey trace is the twelfth trial; below is the difference between the fifth or the twelfth trials and the first trial. Right, corresponding Na^+^ ΔF/F_0_ signals at a site of the AIS associated with the sequential trials on the left; below is the corresponding differences with respect to the first trial. *B*, left, the black trace reports a somatic AP without light exposure ∼20 minutes after establishing whole cell. The grey traces report the somatic AP during 10 consecutive light exposures of 8 ms, with 1 minute interval between two consecutive recordings. The somatic V_m_ was filtered at 10 kHz and recorded at 20 kHz. Right, AP on the left without light exposure is superimposed to the 10^th^ AP with light exposure. The two APs have essentially the same amplitude while the repolarisation of the 10^th^ AP with light exposure is slightly slower. The kinetics of the AP is quantified by the AP amplitude (indicated with *Amp*), calculated as the ΔV_m_ between the beginning and the peak of the AP, and by the half duration (indicated with *D*_*½*_) calculated as the time interval at which V_m_ is above half of its peak value. *C*, left, mean ± SD (N = 6 cells) of *Amp* normalized to the value of the AP before light exposures (*Amp(0)*) indicating that the AP amplitude does not change during the first 10 light exposures. Right, mean ± SD (N = 6 cells) of *D*_*½*_ normalized to the value of the AP before light exposures (*D*_*½*_*(0)*) indicating that the duration of the AP tends to increase during the sequence of light exposures. This increase is however of ∼5% during the first 10 light exposures and its statistical significance is weak (p∼0.05 in the paired t-test calculated between *D*_*½*_*(0)*and *D*_*½*_*(10)*). This increase corresponds to maximum 2-3 samples (100-150 *µ*s), i.e. within the sampling limitations of the patch clamp recording.

### Quantification of the spatial profile of Na^+^ concentration

The spatial resolution of the recordings described above allows reconstructing the spatial profile of the Na^+^ signal along the AIS, associated with an AP. From the representative cell of Fig. 3*A*, we report in Fig. 3*B* the ΔF/F_0_ signal associated with an AP at 6 different sites indicated by their respective distances from the edge of the soma. As shown in Fig.3*B*, the ΔF/F_0_ signals along the AIS have distinct amplitude and kinetics at the different sites. To quantify these ΔF/F_0_ signals in terms of Na^+^ transients, we calibrated the ING-2 fluorescence against the Na^+^ concentration ([Na^+^]) as shown in Fig. 3*C* and we determined that a ΔF/F_0_ transient of 1%, from 10 mM, which is the [Na^+^] in the intracellular solution, corresponds to a 0.175 mM [Na^+^] change (Δ[Na^+^]). With this information, we could reconstruct the spatial profile of the ΔF/F_0_ peak along the AIS and convert it into a Δ[Na^+^] peak profile as shown in the colour coded representation in Fig. 2*D*. The largest Δ[Na^+^] peaks associated with an AP were observed at distances between 15 *µ*m and 25 *µ*m, consistently with the results obtained in a study using another indicator and a slower acquisition rate (Baranauskas *et al.* 2013). The same result was obtained in N = 6 cells (Fig. 3*E*) where this analysis was performed. In these cells, the estimated Δ[Na^+^] peaks were mostly between 1 mM and 1.5 mM, significantly larger (p<0.01, paired t-test) than those closer to the soma (at distances between 5 *µ*m and 10 *µ*m from the soma) and those more distal from the soma (between 30 *µ*m and 35 *µ*m from the soma). Finally, we unambiguously assessed that the ΔF/F_0_ transients, associated with an AP in the AIS, were entirely due to Na^+^ influx via VGNCs by inhibiting APs with extracellular application of 1 *µ*M TTX (Fig. 4*A*) or of 2.5 *µ*M HwTx-IV (Fig. 4*B*), inhibiting the TTX-sensitive VGNCs (Xiao *et al.* 2011). Application of these Na^+^ channels blockers inhibited both the AP and the Na^+^ transients, a result consistently observed in 5 cells tested for each compound (Fig. 4*C*). In summary, these measurements represent a general improvement from previous estimates of the Na^+^ influx in the AIS, associated with APs, using the indicator ANG-2 (Miyazaki & Ross 2015; Miyazaki *et al.* 2019) or the less sensitive indicator SBFI (Kole *et al.* 2008; Fleidervish *et al.* 2010; Baranauskas *et al.* 2013; Scott *et al.* 2014; Katz *et al.* 2018). But the significant and critical improvement of the present measurements is that they were achieved at the unprecedented temporal resolution of 10 kHz. A caption of 8 frames during the rising phase of the AP, in a representative experiment, is illustrated in Fig. 5 using a colour scale. This illustration allows appreciating the temporal resolution of the recording that opens the gate to the reconstruction of the Na^+^ current.

**Figure 3.**
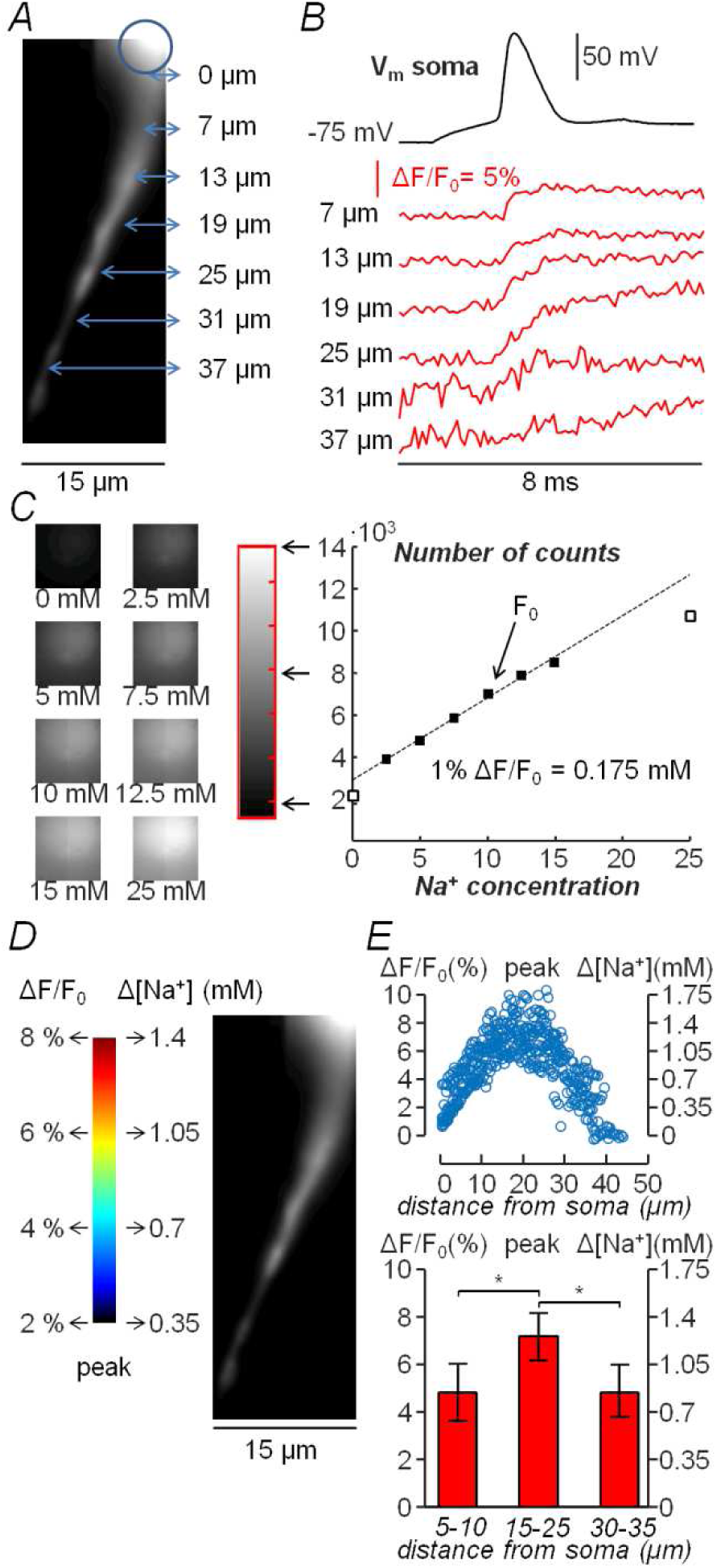
Spatial profile of Na^+^ concentration changes associated with an AP. *A*, AIS of a L5 pyramidal neuron in the recording position with arrows indicating positions at different distance from the soma, calculated from the surface of the cell body in focus (circle). *B*, somatic AP elicited by a 5 ms current step (top, black trace) and ING-2 ΔF/F_0_ associated transients in the six regions indicated in panel *A* (bottom, red traces). Optical traces are from averages of 4 trials. *C*, ING-2 fluorescence images at different Na^+^ concentrations; the grey scale illustrates the conversion between the number of counts and the Na^+^ concentration. Right, plot of the number of counts versus the Na^+^ concentration and linear fit between 2.5 mM and 15 mM Na^+^ concentration. *D*, colour-coded representation of the ΔF/F_0_ peaks along the AIS; the ΔF/F_0_ signal is calibrated in terms of Na^+^ concentration change. *E*, top, plot of the ΔF/F_0_ peak also calibrated as Na^+^ concentration change against the distance from the soma from N = 6 cells. Bottom, from 6 cells above, mean ± SD of the ΔF/F_0_ peak also calibrated as Na^+^ concentration change at 5-10 *µ*m distances from the soma, at 15-25 *µ*m distances from the soma and at 30-35 *µ*m distances from the soma. “*” indicates that the signals are significantly larger at 15-25 *µ*m distances from the soma (p < 0.01, paired t-test). Data are from averages of 4 trials.

**Figure 4.**
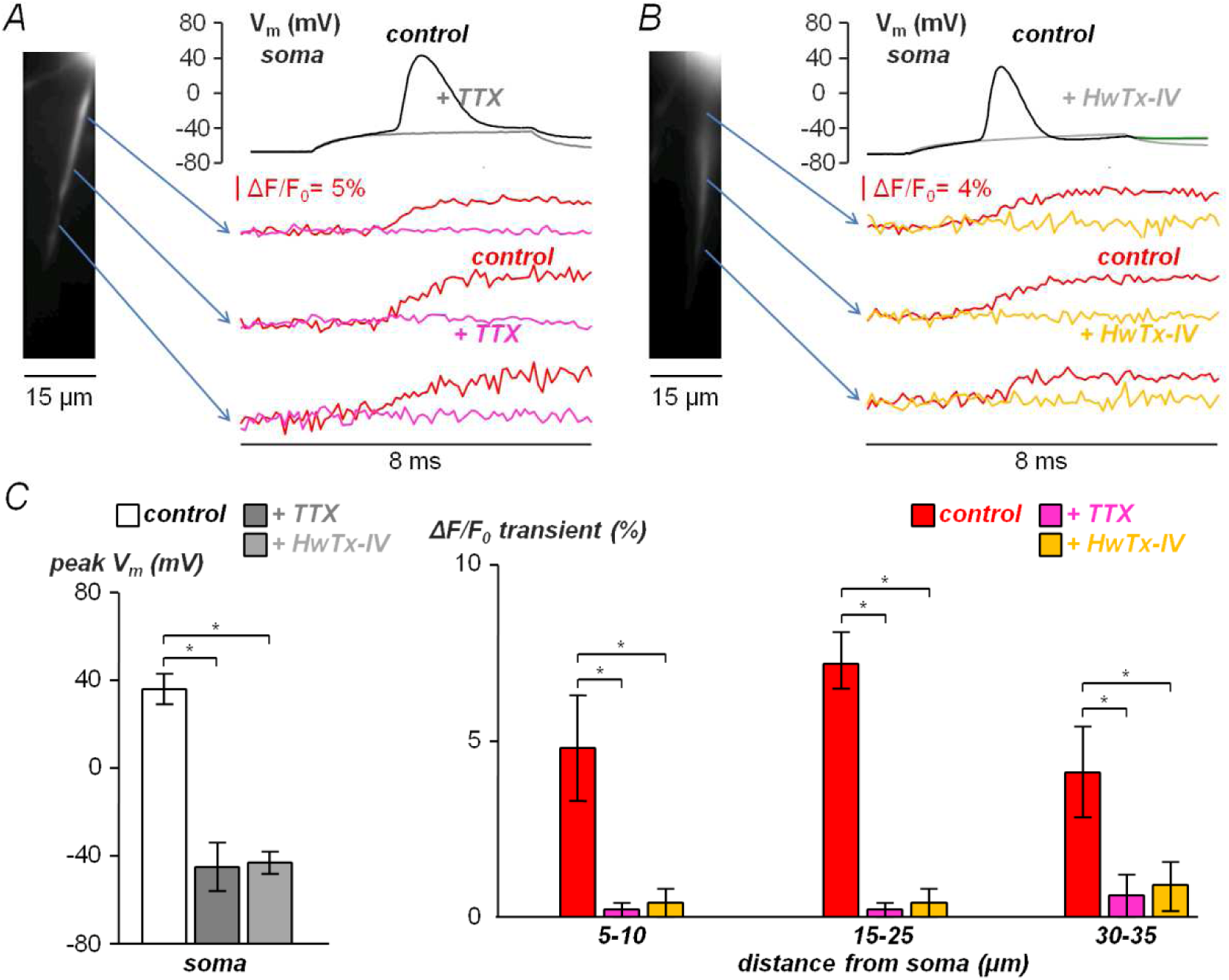
Block of Na^+^ ΔF/F_0_ signals by TTX or by the TTX-sensitive VGNC blocker HwTx-IV. *A*, AP evoked (top-right) in the depicted cell (left) and corresponding ΔF/F_0_ signals in the three positions indicated by the arrows. The AP recorded in control conditions (black trace) is inhibited by local perfusion with a solution containing 1 *µ*M TTX (dark grey trace) and the axonal ΔF/F_0_ transients in control conditions (red traces) are also inhibited (purple traces). *B*, AP evoked (top-right) in another depicted cell (left) and corresponding ΔF/F_0_ signals in the three positions indicated by the arrows. The AP recorded in control conditions (black trace) is inhibited by local perfusion with a solution containing 2.5 *µ*M HwTx-IV (light grey trace) and the axonal ΔF/F_0_ transients in control conditions (red traces) are also inhibited (yellow traces). *C*, left, mean ± SD of the peak V_m_ in control conditions (36 ± 7 mV, N = 10 cells, white columns), and after local application of 1 *µ*M TTX (−47 ± 11 mV, N = 5 cells, dark grey columns) or after local application of 2.5 *µ*M HwTx-IV (−45 ± 5 mV, N = 5 cells, light grey columns). Right, corresponding mean ± SD of the ΔF/F_0_ transients in sampling axonal positions at 5-10 *µ*m, 15-25 *µ*m and 30-35 *µ*m from the soma; red columns are in control conditions (4.8 ± 1.5%, 7.2 ± 0.8% and 4.1±1.3% respectively, N = 10 cells); purple columns are in the presence of TTX (0.2 ± 0.2%, 0.2 ± 0.2%, and 0.6 ± 0.6% respectively, N= 5 cells); yellow columns are in the presence of HwTx-IV (0.4 ± 0.4%, 0.4 ± 0.4%, and 0.9 ± 0.7% respectively, N= 5 cells). “*” indicates significant inhibition (p < 0.01, paired t-test applied to the peaks in the presence of the blockers and the respective peaks in control conditions). VGNC blockers were locally applied by gentle pressure using a pipette (∼10 *µ*m tip) positioned near the AIS. All optical traces are from averages of 4 trials.

**Figure 5.**
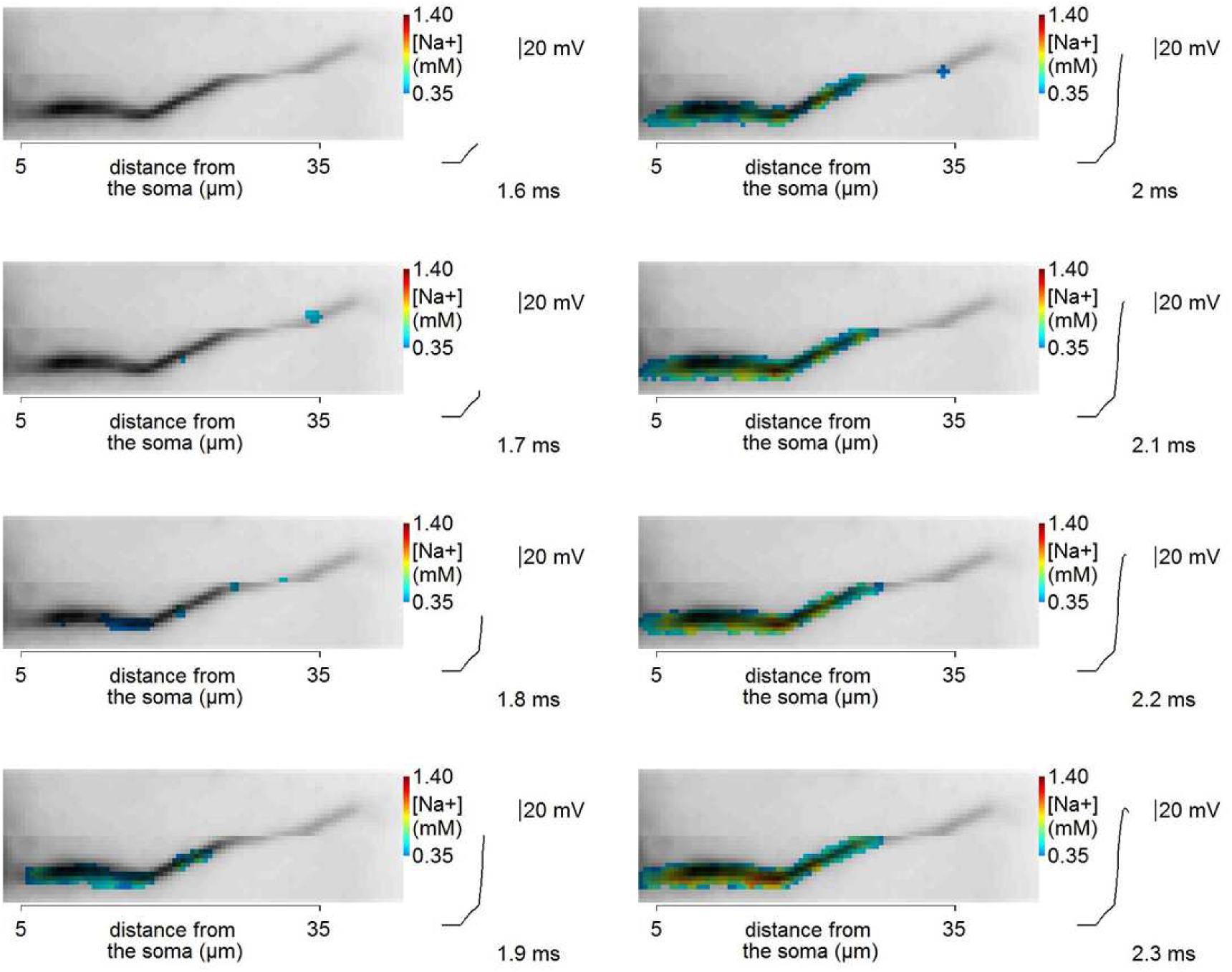
Caption of Na^+^ concentration transient during the AP rising. AIS of a L5 pyramidal neuron and [Na^+^] change during the rise time of an AP depicted in the colour scale indicated. The 8 frames correspond to the instants of the AP rising phase, reported on the right, captured every 100 µs as indicated. Frames are obtained from averages of 4 recordings.

### Determination of the physiological Na^+^ current

Whereas the fluorescence transient of a Ca^2+^ indicator depends on the Ca^2+^ influx, but also on the kinetics of sequestration of Ca^2+^ by endogenous mechanisms (Jaafari *et al.* 2014; Jaafari *et al.* 2015; Ait Ouares *et al.* 2016), the fluorescent transient of a Na^+^ indicator depends instead on the Na^+^ influx and also on the longitudinal Na^+^ diffusion along the axon (Zylbertal *et al.* 2017), while extrusion mechanisms provided by Na^+^ pumps are negligible during the AP (Fleidervish *et al.* 2010). Thus, we initially estimated the small contribution of Na^+^ longitudinal diffusion to the Δ[Na^+^] signal and eventually subtracted it to obtain the Na^+^ influx using the procedure described in the Methods. Specifically, we applied a mask to the fluorescence image (Fig. 6*A*) to estimate the axon diameter d(*x*) at each position and to convert the Δ[Na^+^] was first into the transient of ion density (Δ[N_ions_]) (Fig. 6*B*). We then calculated the compensation factor ([Na^+^]_diff_) and we corrected the original signal (Fig. 6*C*) to take into account diffusion. Finally, we evaluated the contribution of this compensation in the estimate of the Na^+^ current by fitting the compensated signals and the compensating curves with single sigmoid functions and by calculating the time derivative in different regions (Fig. 6*D*). Notably, in N = 27 cells in which the compensation was applied, the peak of the compensating curve was consistently < 5% of the peak of the compensated signal in all regions, indicating that diffusion correction contributes marginally to the estimate of the Na^+^ current in these experiments.

**Figure 6.**
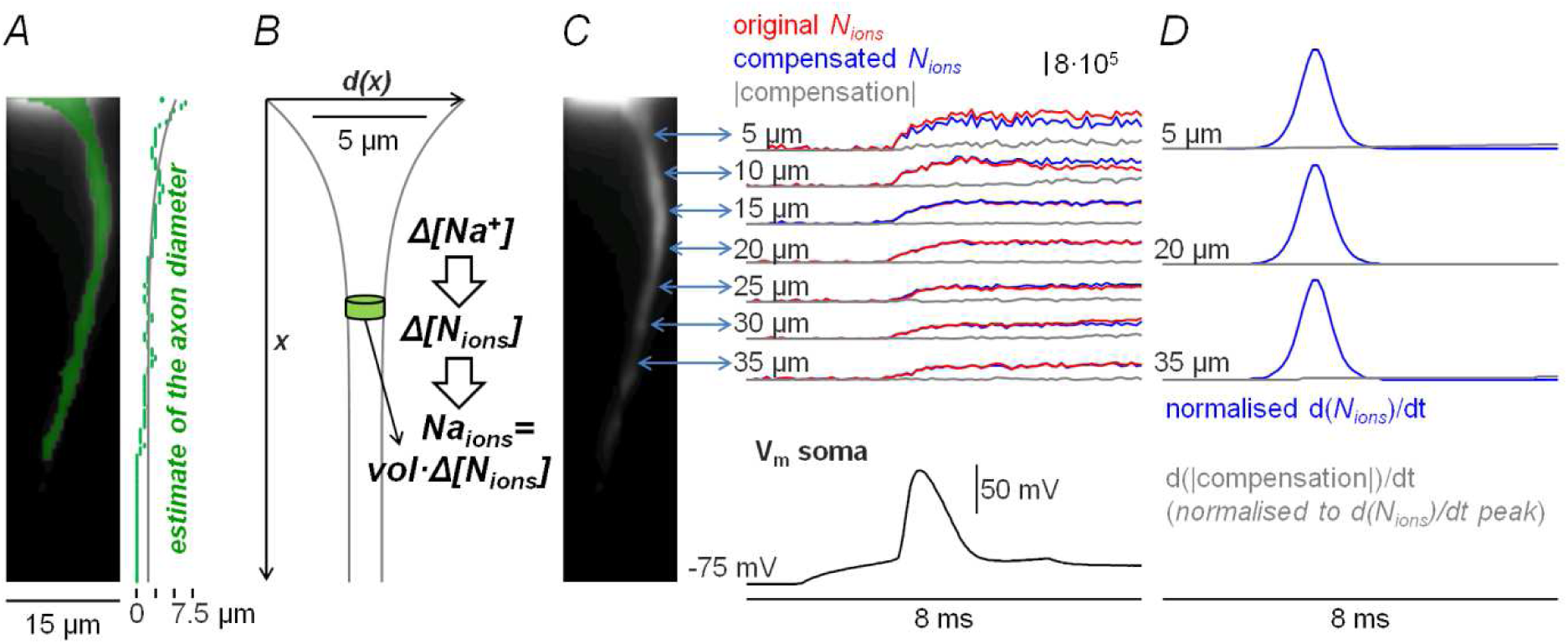
Compensation for longitudinal diffusion. *A*, AIS of a L5 pyramidal neuron in the recording position with a mask (green) indicating the pixels above a certain intensity used to estimate the axon diameter. An equation *d(x)* = *A·x*^*B*^ + *C* was fitted against the *x* axis to obtain a smooth realistic profile. *B*, from the diameter along the *x* axis in panel *A*, estimate of the volume (*vol*) of the disks along the AIS used to convert Δ[Na^+^] into *N*_*ions*_. *C*, the *N*_*ions*_ signal associated with an AP from seven sites at indicated distance from soma. Red traces are the original signals. Blue traces are the signals after compensation for lateral diffusion, using the procedure described in the Methods. Grey traces are the absolute compensating factors (|compensation|). Notice that the ΔN_ions_ signal after compensation is smaller than the original ΔN_ions_ signal at the site nearest to the soma, indicating Na^+^ diffusion in the direction of the soma. *D*, after fitting both the compensated *N*_*ions*_ signals and the compensating factors with sigmoid functions at 5 *µ*m, 20 *µ*m and 35 *µ*m distance from the soma, the time derivatives of the fits normalized to maxima of the d(*N*_*ions*_)/dt signals indicating only small contributions of diffusion to the slope of *N*_*ions*_ signals. Optical traces are from averages of 4 trials.

After correcting the Δ[Na^+^] signals along the positions *x* of the AIS as described above, we converted them into charge surface densities (δQ) calibrated in pC/*µ*m^2^ as illustrated in Fig.7A and as described in the Methods. Notably, the calculated δQ signal is proportional to the radius of the axon which was estimated, in each cell, by a fit of the longitudinal profile of the AIS (see Methods) and it varied from 0.9 to 1.2 *µ*m at 5-10 *µ*m distance from the soma, and from 0.7 to 0.9 *µ*m at 30-35 *µ*m distance from the soma. In this example (Fig. 7*B*), the kinetics of the δQ signals was initially characterised by a first subthreshold component, above the noise of the initial samples, beginning after the pulse of somatic current injection but before the somatic AP. The δQ starting at the beginning of the somatic AP was then characterised by a steep (fast) increase followed by a slower increase as indicated by the two arrows. Interestingly, a biphasic Na^+^ transient in the AIS was already reported by recording SBFI fluorescence at 2 kHz (Baranauskas *et al.* 2013). To reconstruct the Na^+^ current (I_Na_) by calculation of the time derivative, we first fitted the δQ signals with the two-component function reported in the “Signals fit and calculation of the Na^+^ current” section of the Methods. The rationale of this model function is also explained in the Methods and it is illustrated in Fig. 8. The I_Na_ was then obtained for the different sites by calculating the time derivative of the fitted model function (Fig. 7*B*), in which the values of the parameters α, γ, η_3_, ν_2_ and ν_3_ were obtained as described in the Methods. Notably, the suprathreshold current was composed of a fast current peaking at *t* = η_1_ at the beginning of the rising phase of the AP, and of a slow current persisting during the repolarisation of the AP. We assessed the ability of this model function to faithfully reproduce the kinetics of the physiological I_Na_ by comparing the noisy shape of the time derivative of the experimental signal, filtered with a smoothing Savitsky-Golay algorithm, with the shape of the time derivative of the model function fitted to the same experimental signal. The comparison shows that the model function reproduces the kinetics of the time derivative of the original signal, validating its use in the measurement of the Na^+^ current. We next analysed how the I_Na_ signals vary from one cell to another. Fig. 9*A* shows δQ signals from another cell from proximal (10 *µ*m from the soma), medial (20 *µ*m from the soma) and distal (30 *µ*m from the soma) sites of the AIS. In this example, the width of the somatic AP, quantified as the interval during which V_m_ > 0, was wider with respect to the case of Fig.7. The I_Na_ signals obtained for this cell (Fig. 9*B*) have fast components characterised by peak amplitudes similar to those of the previous cell, preceding the peak of the somatic AP by an interval Δ*t*, but longer lasting slow components quantified by the parameter ν_3_. We reconstructed the I_Na_ from different sites of the AIS in N = 15 cells, and in 6 cells a clear subthreshold component of the current above the photon noise was detected. As shown in Fig. 9*C*, the peaks of the fast current at sites located at 5-10 *µ*m from the soma (75 ± 21 pA/*µ*m^2^) were comparable to those measured at sites located at 15-25 *µ*m from the soma (74 ± 29 pA/*µ*m^2^), a result indicating that the smaller Na^+^ transient observed near the soma (see Fig. 9*D*) is possibly due to the smaller surface-to-volume ratio decreasing the [Na^+^] change, although this result depends on our estimate of the axonal radius. In contrast, the peaks of the fast current at sites located at 30-35 *µ*m from the soma (35 ± 13 pA/*µ*m^2^) were significantly smaller (Fig. 9*C*). The peak of the fast currents was consistently observed during the rising phase of the somatic AP, with an interval Δ*t* that was 240 ± 89 *µ*s, 250 ± 82 *µ*s and 253 ± 83 *µ*s at proximal medial and distal sites respectively (Fig. 9*D*). This interval was comparable to the interval between the maximal slope of the AP, (i.e. the time of the peak of the AP time derivative), and of the AP peak (313 ± 157 *µ*s). The timing of the somatic AP, however, does not correspond to the peak of the AP in the AIS since the axonal AP upstroke is steeper than the somatic AP upstroke (Popovic *et al.* 2011; Popovic *et al.* 2015). To evaluate the actual timing of the AP in the AIS, we performed V_m_ imaging recordings to measure simultaneously axonal and somatic APs, using the same protocols of Na^+^ imaging experiments. As shown in the example of Fig. 10*A*, the occurrence of the onset of the axonal and somatic APs is within one camera sample (100 *µ*s), but the peak of the somatic AP is delayed by 1-2 frames from the peak of axonal APs. This delay of 1-2 frames was consistently observed in N = 5 cells tested in this way (Fig. 10*B*). It follows that the peak of the fast current occurs during the rising phase of the AP, before the peak of the AP. We finally tested the correlation between the values of w*t* and the values of the parameter ν_3_ quantifying the duration of the slow current (Fig. 9*E*). We calculated the Pearson linear coefficient and we found that these parameters were significantly negatively correlated (p<0.01) at all sites of the AIS, indicating that the slow component, occurring during the repolarisation phase of the AP, is correlated with its duration. In summary, we provided the first experimental measurement of the physiological Na^+^ current associated with the AP.

**Figure 7.**
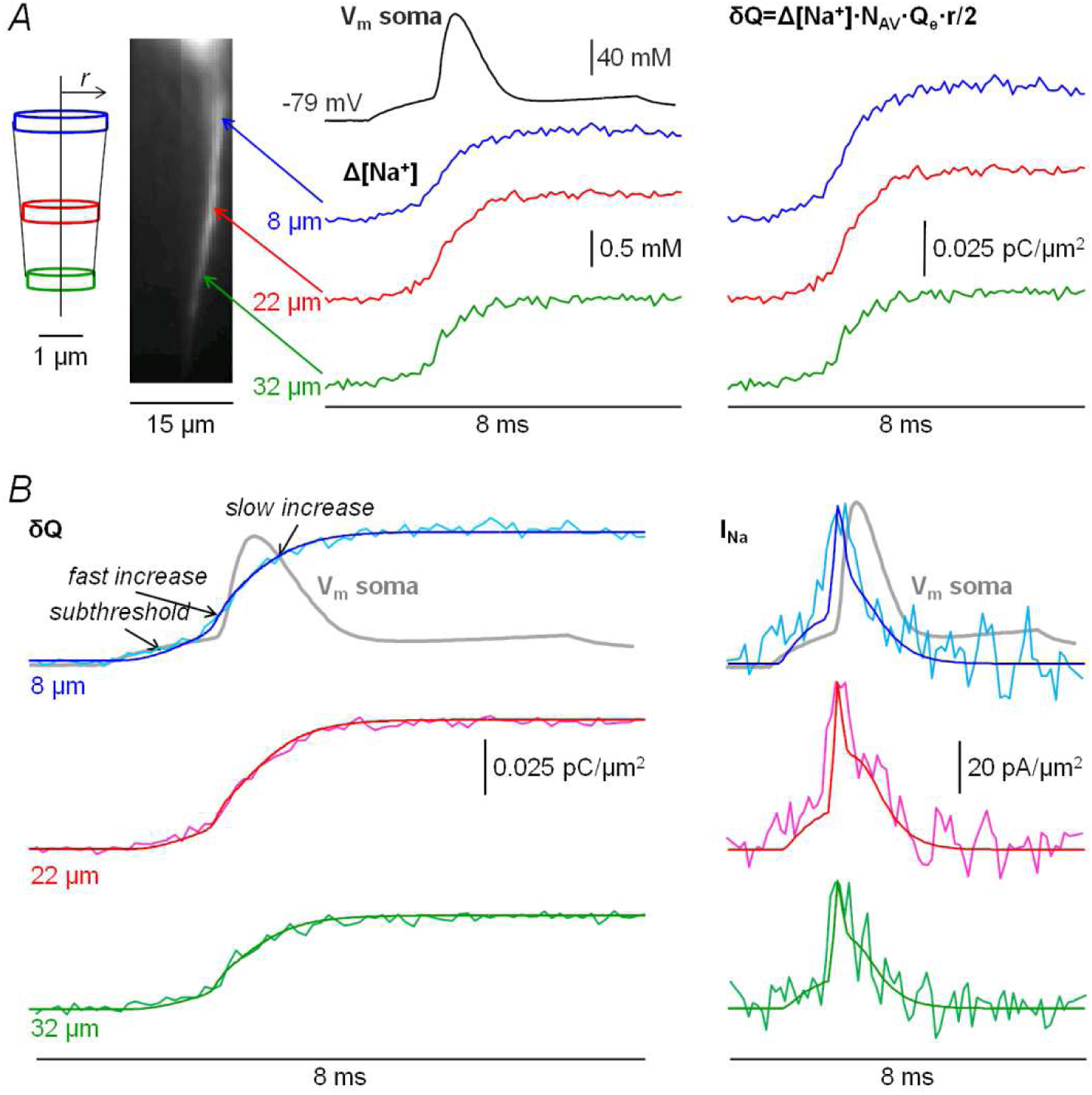
Reconstruction of the Na^+^ current associated with an AP in the AIS. *A*, left, AIS of a L5 pyramidal neuron in the recording position with arrows indicating positions at 8 *µ*m (blue), 22 *µ*m (red) and 32 *µ*m (green) from the soma. The reconstruction of the surface of the AIS is illustrated. Middle, somatic AP elicited by a 5 ms current step (top, black trace) and the Δ[Na^+^] signals from the three sites indicated on the left. Right, the charge density (*δQ*) signals calculated as indicated, with N_AV_ and Q_e_ indicating the Avogadro number and the elementary charge respectively. *B*, left, *δQ* signals in panel *A* fitted with the model function. The arrows point three phases of the *δQ* increase corresponding to a subthreshold component (between the beginning of the somatic current injection and the AP onset), a fast increase at the onset of the AP and a slow increase during the repolarisation of the AP. Right, the Na^+^ current (I_Na_) at the different sites obtained by calculating the time derivatives of the fits. Superimposed are the time derivatives of the *δQ* signals filtered by a Savitzky-Golay smoothing algorithms with a span of 10 points. The somatic AP (grey trace) is superimposed to the top *δQ* and I_Na_ traces. Data are from averages of 4 trials.

**Figure 8.**
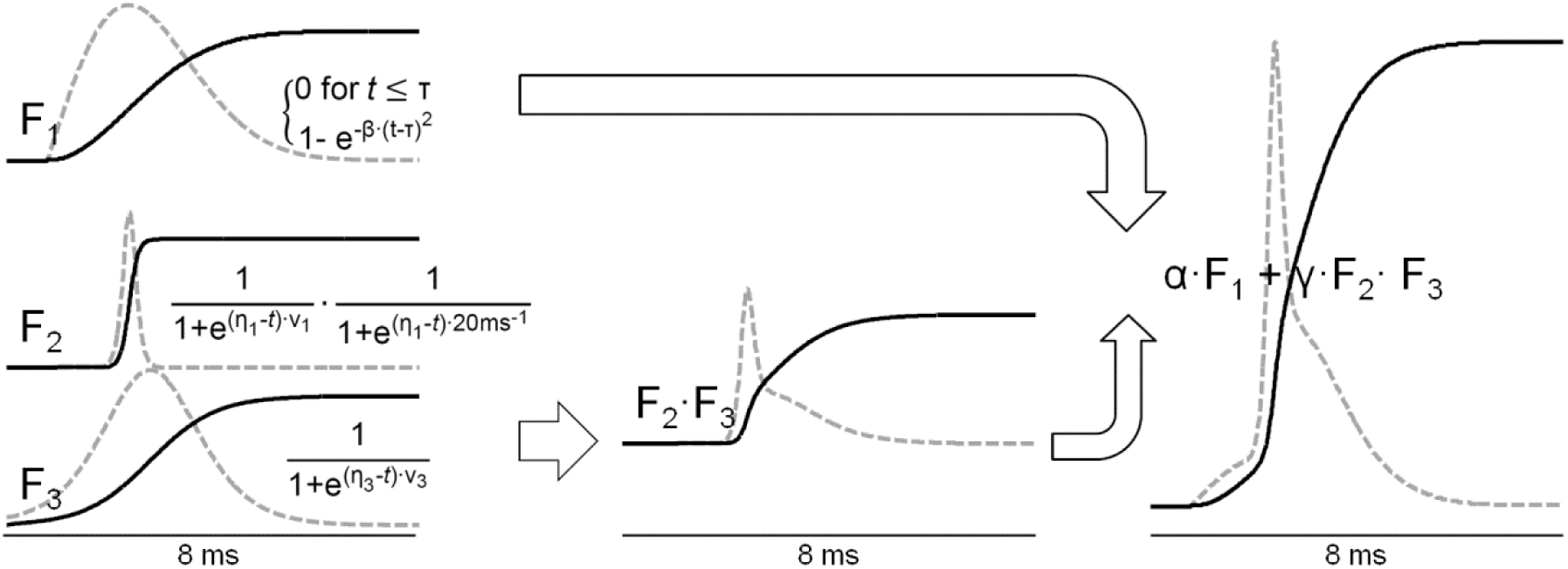
Rationale of the model function used to fit Na^+^ transients. Left, the three functions (black traces) composing the model function used to fit the Na^+^ transient. The normalized time derivatives are superimposed (grey dotted traces). F_1_ is the function used to match the subthreshold component with dF_1_/d*t* = 0 before the injection of the somatic current (starting at t = τ) and slowly increasing after the beginning of the somatic current (t > τ). F_2_ is the function used to match the fast phase of the suprathreshold component with dF_2_/d*t* = 0 before the beginning of the AP and 1 ms after the AP peak. F_3_ is the function used to match the slow phase of the suprathreshold component persisting during the AP repolarisation. Middle, the product F_2_·F_3_ resulting in the suprathreshold component with d(F_2_·F_3_)/d*t* = 0 before the beginning of the AP, but with d(F_2_·F_3_)/d*t* > 0 during the AP repolarisation. Right, the final model function α·F_1_ + γ·F_2_·F_3_ reproducing the composite kinetics of the Na^+^ transient and allowing the reconstruction of the I_Na_ when fitting the free parameters on the δQ signal.

**Figure 9.**
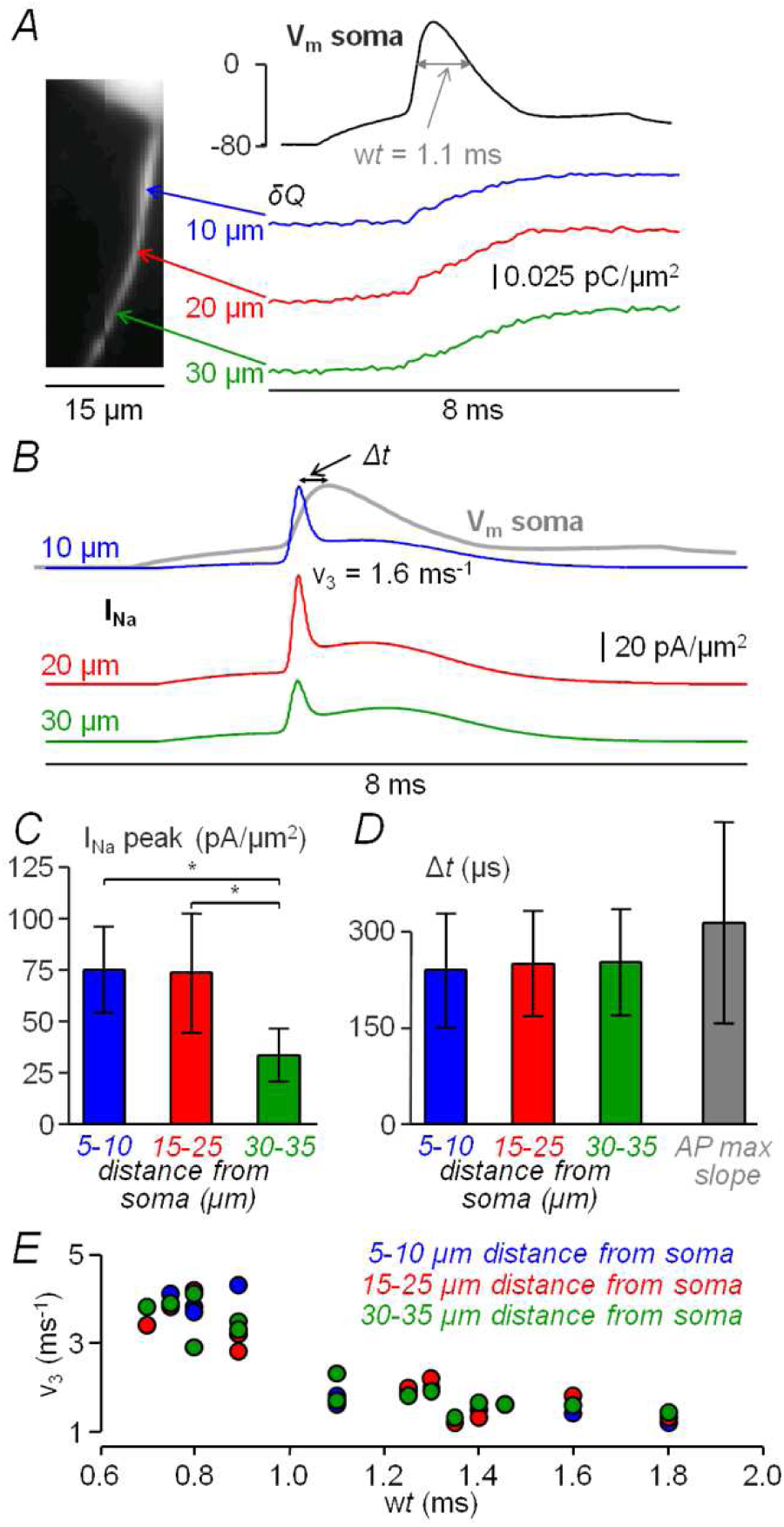
Analysis of I_Na_ currents. *A*, left, AIS of L5 pyramidal neuron in recording position with arrows indicating positions at 10 *µ*m (blue), 20 *µ*m (red) and 30 *µ*m (green) from the soma. Right, somatic AP elicited by a 5 ms current step (top, black trace) and charge density (*δQ*) signals from the three sites on the left. The grey double arrow indicates the width of the somatic AP (*wt*), defined as the V_m_ interval above 0 mV. *B*, Na^+^ currents (I_Na_) calculated from the signals in panel *A*. The somatic AP (grey trace) is superimposed to the top I_Na_ trace. Δ*t* is the interval between the peak of the I_Na_ and the peak of the somatic AP. The value of the fitting ν_3_ parameter shaping the slow I_Na_ component is reported for the top I_Na_. Data are from averages of 4 trials. *C*, mean ± S.D. (N = 15 cells) of the fast peak of I_Na_ from N = 15 cells from sites at 5-10 *µ*m (blue column), 15-25 *µ*m (red column) and 30-35 *µ*m (green column) from the soma. “*” indicates that peaks at 30-35 *µ*m distance were significantly smaller than those at 5-10 *µ*m and 15-25 *µ*m distance. *D*, mean ± S.D. (N = 15 cells) of Δ*t* from sites at 5-10 *µ*m, 15-25 *µ*m and 30-35 *µ*m from the soma. The grey column on the right is the interval between the maximum AP slope and the somatic AP peak. *E*, from N = 15 cells, plot of the values of the fitted ν_3_ parameter of the model function against the AP interval w*t*. At all distances, the two variables were negatively correlated (p< 0.01, Pearson coefficient −0.85, −0.83 and −0.91 for proximal, medial and distal sites respectively). Data are from averages of 3-6 trials.

**Figure 10.**
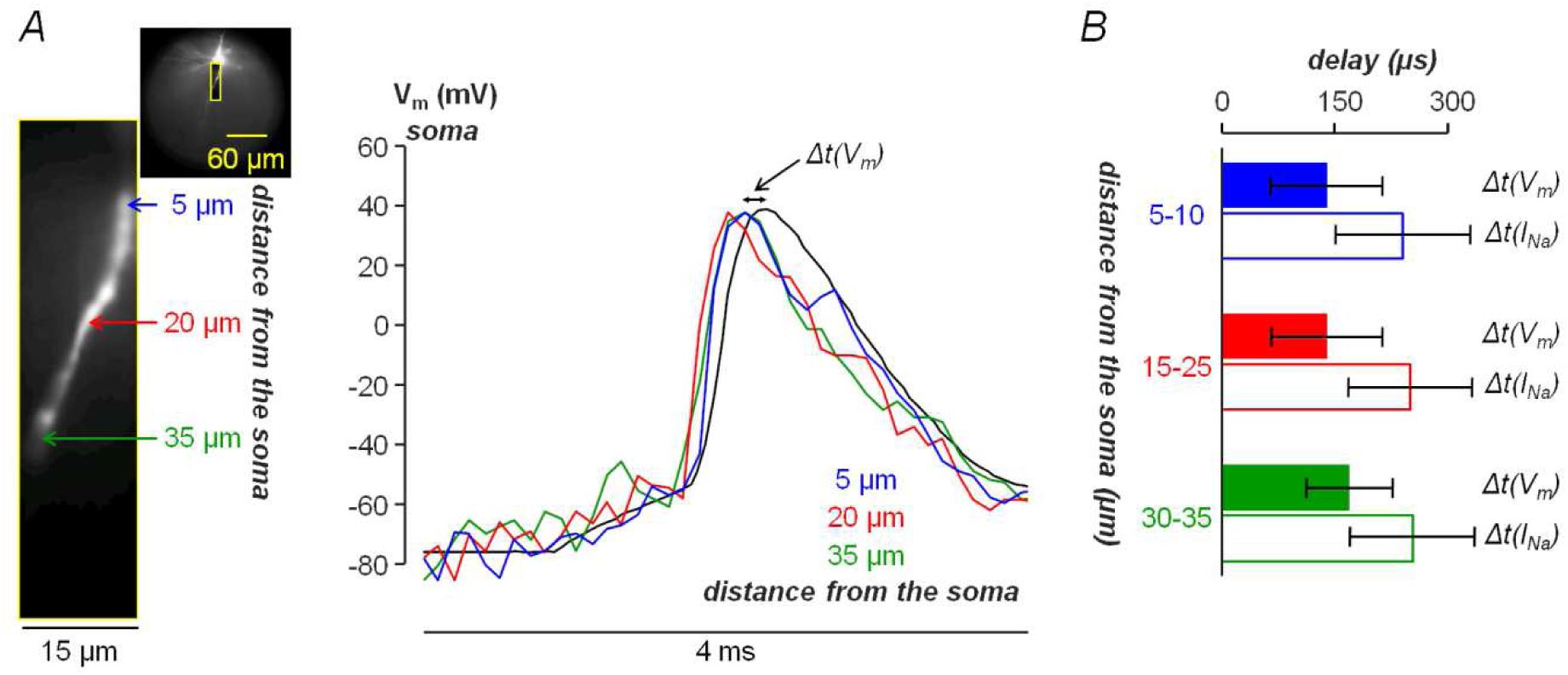
AP in the AIS measured using V_m_ imaging. *A*, left, AIS of a L5 pyramidal neuron filled with the voltage sensitive dye JPW1114 in the recording position with three sites indicated at 5, 20 and 35 *µ*m from the soma. Right, somatic AP superimposed to the axonal AP at 5 *µ*m (blue trace), 20 *µ*m (red trace) and 35 *µ*m (green trace) distance from the soma. The fluorescent transients associated with the AP are normalised to the somatic AP. The delay between the axonal AP peak and the somatic AP peak (*Δt(V*_*m*_)) is illustrated with a double arrow. All traces are from averages of 4 trials. *B*, colour-filled columns are mean ± SD (N = 5 cells) of the delay between the axonal AP peak and the somatic AP peak (*Δt(V*_*m*_)) at 5-10 *µ*m from the soma (140 ± 74 *µ*s, blue columns), at 15- 25 *µ*m from the soma (140 ± 74 *µ*s, red columns) and at 30-35 *µ*m from the soma (170 ± 57 *µ*s, green columns). Colour-outlined columns are mean ± SD (N = 15 cells) of the delay between the reconstructed axonal I_Na_ peak (from Na^+^ imaging experiments) and the somatic AP peak (*Δt(I*_*Na*_)) also reported in Fig. 9*D* at 5-10 *µ*m from the soma (blue columns), at 15-25 *µ*m from the soma (red columns) and at 30-35 *µ*m from the soma (green columns).

### Comparison between optically measured and simulated Na^+^ currents

Several detailed models of L5 pyramidal neurons have been built in order to predict the Na^+^ currents underlying AP generation and propagation (Kole *et al.* 2008; Hu *et al.* 2009; Fleidervish *et al.* 2010; Hallermann *et al.* 2012; Baranauskas *et al.* 2013; Cohen *et al.* 2020). Hence, we next compared our experimentally measured I_Na_ signals with the Na^+^ currents obtained by running simulations with two NEURON published models from Fleidervish et al. (2012) (available in the database https://senselab.med.yale.edu/ModelDB/ShowModel?model=136715) and from Hallermann et al. (2012) (available in the database https://senselab.med.yale.edu/ModelDB/ShowModel?model=144526). Fig.11*A* shows the I_Na_ signals already reported in Fig. 7*B* in proximal, medial and distal sites of the AIS and the currents from the two models in equivalent AIS positions. The Na^+^ currents from the Fleidervish et al. model, associated with an AP having similar kinetics with respect to the measured one, had fast components which were nearly half of the experimental ones and slow components of similar duration with respect to the experimental ones. In contrast, the Na^+^ currents from the Hallermann et al. model, associated with a sharper AP with respect to the measured one, had larger and sharper fast components with respect to the experimental ones and negligible slow components with respect to the experimental ones. Thus, to obtain simulated APs and Na^+^ currents more similar to the experimental ones, we modified the activation and inactivation functions of the two VGNCs in the Hallermann et al. model as described in the Methods. Fig. 11*B* shows the APs and the medial I_Na_ signals of the cell in Fig. 7 (cell 1) and of the cell in Fig. 9 (cell 2) superimposed to display the longer width of the AP in cell 1 and the consequent longer lasting slow I_Na_ component. The results of two simulations, where the density of K^+^ channels was the same of that in the original model in simulation 1, and 66% of that in the original model in simulation 2, are reported in Fig. 11*C*. The two Na^+^ currents were processed with a 100 *µ*s moving average filter to be consistent with the experimental acquisition rate. Using the modified VGNC functions, the filtered simulated Na^+^ currents had amplitude and kinetics similar to the experimental I_Na_ signals with a longer slow component corresponding to the wider AP. In agreement with the experimental case, quantified by the correlation between the parameter ν_3_ and the width of the AP (Fig. 9*E*), the slow component of the Na^+^ current is lasting longer in the simulation producing a wider AP (Fig. 11*C*). Notably, we also estimated the metabolic cost in the modified model by calculating the numbers of ATP molecules required to pump the Na^+^ out. For simulations 1 and 2, these were 1.87·10^7^ and 1.46·10^7^ respectively in the AIS, and 2.02·10^8^ and 1.77·10^8^ respectively in the whole cell. These numbers were close to those in the original model, i.e. 2.20·10^7^ in the AIS and 1.95·10^8^ in the whole cell, indicating that our Na^+^ channel modifications produce only marginal changes in the metabolic cost of the AP generation. Whereas this component is due to the slow inactivating channels, the fast component is due to fast inactivating channels that recover in several milliseconds (Martina & Jonas, 1997), causing a decrease in the amplitude of a second AP occurring at a short time interval from the first one. This relatively long recovery period is incorporated in the Hallermann et al. model and the phenomenon of decrease of the second AP amplitude is reproduced by the computer simulation reported in Fig. 12*A*, where two consecutive pulses of 4.5 ms duration and 6 ms interval between the beginnings of the two pulses are used as stimulation protocol. In the cell reported in Fig. 12*B*, we used the same stimulation protocol to elicit two APs of different amplitudes and measured the Na^+^ fluorescence transient, allowing a comparison between the simulated (Fig. 12*C*) and measured (Fig. 12*D*) Na^+^ currents. While a decrease of the fast component of the second Na^+^ current is the consequence of the VGNC functions, the decrease of the second I_Na_ fast component was quantitatively appreciated in the experiment. Interestingly, the wider second AP is again correlated with a longer duration of the second I_Na_ slow component also within this protocol.

**Figure 11.**
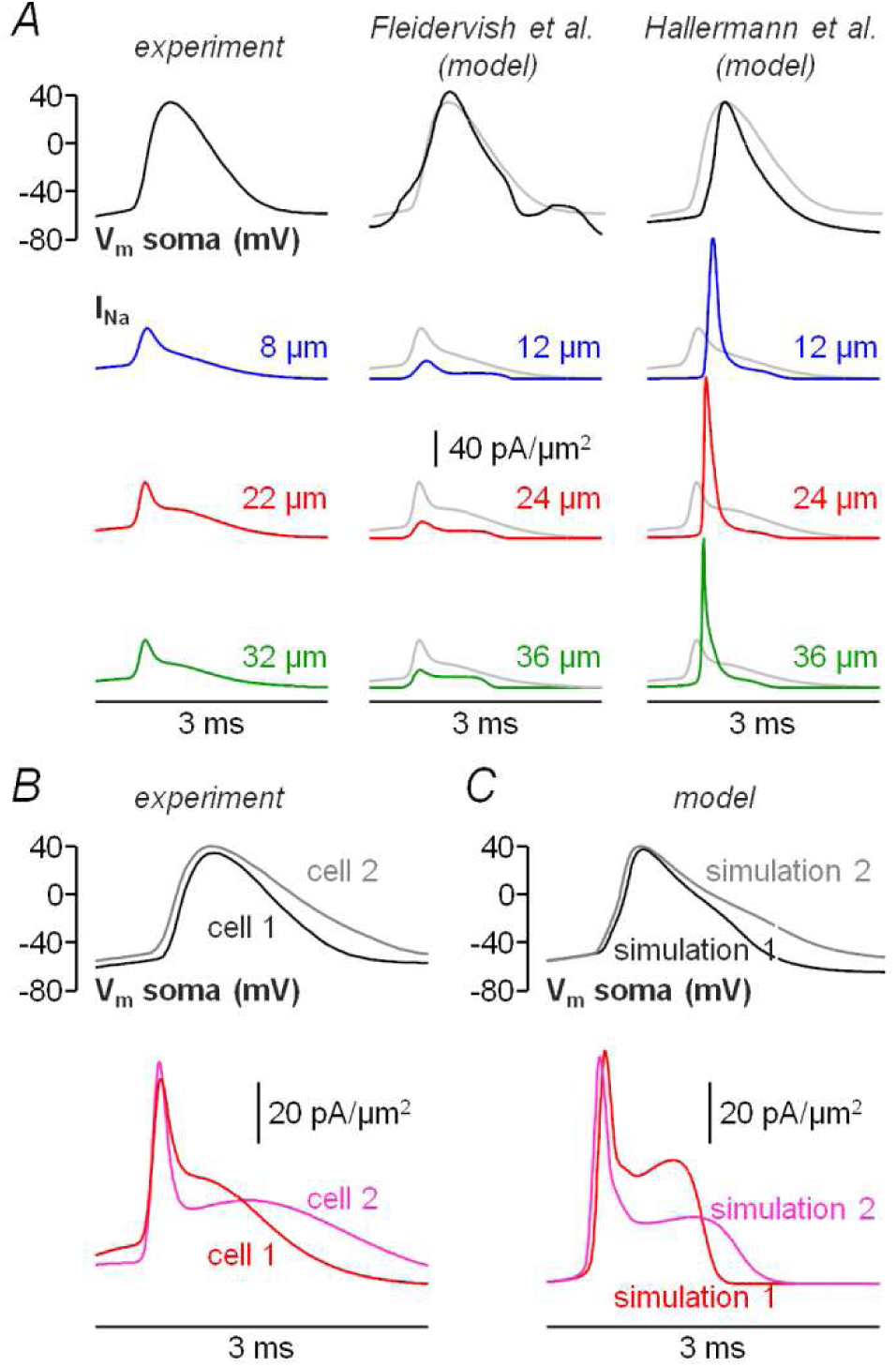
Comparison between optically measured and simulated Na^+^ currents. *A*, left, the AP and the three I_Na_ signals from the cell reported in Fig. 7. Middle and right, the APs and the Na^+^ currents at 12 *µ*m, 24 *µ*m and at 36 *µ*m distances from the soma from computer simulations using the model reported in Fleidervish et al. (2010), and from the model reported in Hallermann et al. (2012). The experimental I_Na_ signals are superimposed (grey traces). *B*, APs and medial axonal I_Na_ signals from the two cells reported in Fig. 7 and Fig. 9 exhibiting different width of the AP and different length of the slow current component. *C*, APs and associated Na^+^ currents from two computer simulations using the Hallermann et al. model where the activation and the inactivation functions of the two VGNCs were modified as described in the Methods, and the density of the K^+^ channels was varied in order to modulate the width of the AP. Na^+^ currents were processed with a 100 *µ*s moving average filter to be consistent with the experimental currents. The duration of the slow current component is longer when the AP is wider, in agreement with the experimental analysis reported in Fig. 9*E*.

**Figure 12.**
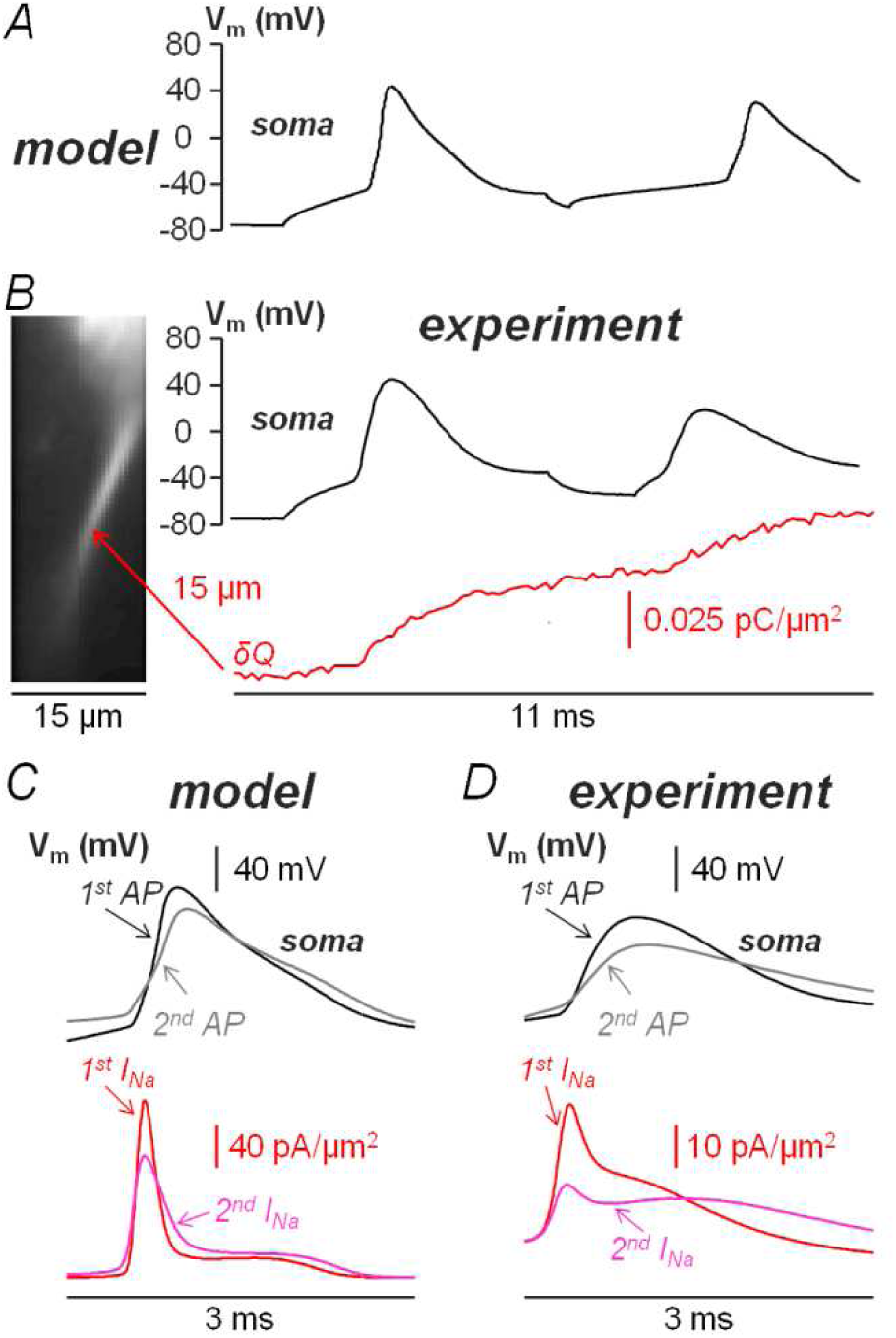
Na^+^ currents associated with two consecutive APs at short time interval. *A*, using the modified model of Fig. 11*C* corresponding to the wider AP, two APs from a computer simulation with 4.5 ms duration pulses and 6 ms interval between the beginnings of the two pulses. *B*, from the cell on the left, two APs elicited by the same stimulation protocol (two pulses with 4.5 ms duration pulses and 6 ms interval) used for the computer simulation in panel *A*. The measured axonal δQ signal at 15 *µ*m distance from the soma is reported below (red trace). Data are form averages of 4 trials. *C*, top, the two APs of the simulation in panel *A* superimposed. Bottom, the corresponding Na^+^ currents processed with a 100 *µ*s moving average filter. The second smaller AP is associated with a smaller fast component of the current. *D*, top, the two APs of the experiment in panel *B* superimposed. Bottom, the corresponding measured Na^+^ currents. As in the case of the computer simulation, the fast component of the I_Na_ signal associated with the second AP is smaller.

In summary, the present approach can be used to build more realistic models of the AIS, but this comparison between experimental and simulated currents also demonstrates that our novel method is capable of experimentally measuring, cell by cell, the behaviour of both fast inactivating and slow inactivating VGNCs at the site of AP origin, without the need of introducing a model.

## DISCUSSION

Experimental approaches for the investigation of Na^+^ influx in the intact AIS during AP generation include the voltage-clamp patch electrode technique (Yue *et al.* 2005; Carter & Bean, 2009; Katz *et al.* 2018) and the optical recording of fluorescence from Na^+^ indicators (Kole *et al.* 2008; Fleidervish *et al.* 2010; Baranauskas *et al.* 2013; Miyazaki & Ross, 2015; Katz *et al.* 2018; Miyazaki *et al.* 2019). The first approach allows recording fast Na^+^ currents triggered by artificial somatic injections of voltage waveforms mimicking an AP, with no information on the spatial organisation of the current and limitations due to space-clamp problems (Bar-Yehuda & Korngreen, 2008). The second approach allows monitoring the spatial profile of Na^+^ transients during physiological APs, but with a temporal resolution insufficient to track the kinetics of Na^+^ currents. Here we report the critical improvement in the Na^+^ imaging technique overcoming the above limitation, permitting the reconstruction of Na^+^ currents associated with physiological APs at different positions of the AIS. Thanks to this achievement, it is now possible to reconstruct Na^+^ currents underlying the AP generation in many neuronal types. We developed this innovative approach in L5 neocortical pyramidal neurons characterised by visually distinguishable axons and we measured fluorescence at distances ranging from 5 to 35 *µ*m from the soma. In this system, we were able to resolve Na^+^ fluorescence transients at 10 kHz with a pixel resolution of 500 nm in single trials. Notably, the remarkable SNR allowed detecting, in a proportion of cells, a subthreshold component of the Na^+^ influx occurring during the somatic current injection but before the onset of the AP. A subthreshold Na^+^ current amplifies the transmission of depolarisation from the soma to the axon and it is therefore expected to contribute to the transduction of postsynapic excitatory inputs from the dendrites into the firing output of the neuron (Goldwyn *et al.* 2019). The principal (suprathreshold) current starting at the onset of the AP was visually characterized by two phases: a first phase with a steep fluorescence increase and a second phase with a longer lasting and weaker fluorescence increase due to a slow Na^+^ current. The first phase leads to a fast current consistently occurring during the rise of the AP that can be classified as *fast inactivating current* (Stuart & Sakmann, 1994). The second phase leads to a slow current occurring during the AP repolarisation and its duration is correlated with the width of the AP. This current can be classified as *persisting slow inactivating current* (Yue *et al.* 2005; Astman *et al.* 2006). The fast current is in its decaying phase at the AP peak when the driving force for Na^+^ is at its minimum, whereas the peak of the slow current occurs at relatively low V_m_, indicating that the channels mediating this current close at more negative V_m_. The two currents were associated with the two VGNCs in a published model (Hallermann *et al.* 2012). By modifying the activation and inactivation functions of these channels, we obtained simulated currents similar to the experimental ones, with marginal changes in the metabolic cost predicted by the original model. Notably, while an acquisition rate of 100 kHz is expected to be necessary to fully reconstruct the kinetics of the Na^+^ current (Popovic *et al.* 2011), the peak of the experimental current obtained at 10 kHz can be still quantitatively appreciated. Thus, our approach allows measuring directly the physiological Na^+^ current associated with an AP in many neuronal types without the need of complex models based on many untested assumptions necessary to set the multiple parameters. This is particularly important since Na^+^ currents are typically mediated by more than one VGNC type with different spatial arrangement and biophysical properties. Specifically, our measurement permits in principle the association of the different current components with their sources along the AIS. For instance, in the case of L5 pyramidal neurons, the Na^+^ current in the AIS is mediated by Nav1.2 and Nav1.6 with different axonal distributions (Hu *et al.* 2009) and electrophysiogical properties (Rush *et al.* 2005). Similarly, our measurement can be used to investigate the physiological modulation of VGNCs by several secondary messengers such as serotonin (Yin *et al.* 2017) or brain-derived neurotrophic factors (Ahn *et al.* 2007), but also the Na^+^ currents in animal models with genetic impairment of VGNCs and related neurological disorders. In particular, critical mutations of Nav1.6 (Wagnon *et al.* 2015) and Nav1.2 (Reynolds *et al.* 2020) are correlated with infant epilepsy, autism and related severe neurological disorders. Interestingly, in L5 pyramidal neurons, Nav1.6 deficient mice are characterized by a reduced persistent Na^+^ current (Katz *et al.* 2018) whereas Nav1.2 deficient mice are characterized by reduced dendritic excitability and synaptic function (Spratt *et al.* 2019).

The present approach can be improved further in the future by increasing the acquisition rate and the number of possible trials within one experiment while preserving the same SNR, which represents the bottom neck limitation of the technique. The state-of-the-art of our novel approach is ultimately depending on the dye ING-2, which has a remarkable sensitivity but also some intrinsic limitations. While the Na^+^ indicator SBFI has been used at 2 mM concentration (Fleidervish *et al.* 2010), 500 *µ*M was the highest tolerated concentration of ING-2 that did not cause a change in the shape of the somatic AP recorded after establishing the whole-cell configuration. Clearly, a more inert and photo-stable indicator with equal sensitivity and quantum efficiency, but tolerated at higher concentrations, would allow larger numbers of trials at higher acquisition rates. This achievement would result in a more accurate sampling of the signal and in a higher number of possible trials, which is 10 at present. Thus, critical improvements of our novel approach are foreseen in the future by using novel Na^+^ indicators.

## Acknowledgements

This work was supported by the *Agence Nationale de la Recherche* through three grants (ANR-18-CE19-0024 - OptChemCom; Labex *Ion Channels Science and Therapeutics:* program number ANR-11-LABX-0015 and National Infrastructure France Life Imaging “Noeud Grenoblois”) and by the *Federation pour la recherché sur le Cerveau* (FRC – Grant *Espoir en tête*, Rotary France). We thank Dr. Marko Popovic for helping us in setting up the preparation and the optical measurements and Prof. Bertrand Fourcade for helping us in developing the strategy for taking into account sodium lateral diffusion.

## REFERENCES

Ahn M, Beacham D, Westenbroek RE, Scheuer T & Catterall WA (2007). Regulation of Na(v)1.2 channels by brain-derived neurotrophic factor, TrkB, and associated Fyn kinase. J Neurosci 27, 11533–11542.

Ait Ouares K, Filipis L, Tzilivaki A, Poirazi P & Canepari M (2019). Two Distinct Sets of Ca2+ and K+ Channels Are Activated at Different Membrane Potentials by the Climbing Fiber Synaptic Potential in Purkinje Neuron Dendrites. J Neurosci 39, 1969–1981.

Ait Ouares K & Canepari M (2020). The origin of physiological local mGluR1 supralinear Ca_2+_ signals in cerebellar Purkinje neurons. J Neurosci 40, 1795–1809.

Ait Ouares K, Jaafari N & Canepari M (2016). A generalised method to estimate the kinetics of fast Ca(2+) currents from Ca(2+) imaging experiments. J Neurosci Methods 268, 66 –77.

Astman N, Gutnick MJ & Fleidervish IA (2006). Persistent sodium current in layer 5 neocortical neurons is primarily generated in the proximal axon. J Neurosci 26, 3465–3473.

Baranauskas G, David Y & Fleidervish IA (2013). Spatial mismatch between the Na+ flux and spike initiation in axon initial segment. Proc. Natl. Acad. Sci USA 110, 4051–4056.

Bar-Yehuda D & Korngreen A (2008). Space-clamp problems when voltage clamping neurons expressing voltage-gated conductances. J Neurophysiol 99, 1127–1136.

Bean BP (2007). The action potential in mammalian central neurons. Nat Rev Neurosci 8, 451–465.

Canepari M, Vogt K & Zecevic D (2008). Combining voltage and calcium imaging from neuronal dendrites. Cell Mol Neurobiol 28, 1079–1093.

Canepari M, Willadt S, Zecevic D & Vogt KE (2010). Imaging Inhibitory Synaptic Potentials Using Voltage Sensitive Dyes. Biophys J 98, 2032–2040.

Carter BC & Bean BP (2009). Sodium entry during action potentials of mammalian neurons: incomplete inactivation and reduced metabolic efficiency in fast-spiking neurons. Neuron 64, 898–909.

Cohen CCH, Popovic MA, Klooster J, Weil MT, Möbius W, Nave KA & Kole MHP (2020). Saltatory Conduction along Myelinated Axons Involves a Periaxonal Nanocircuit. Cell 180, 311–322.

Crank J (1975). The mathematics of diffusion. (Oxford: Clarendon Press).

Filipis L, Ait Ouares K, Moreau P, Tanese D, Zampini V, Latini A, Bleau C, Bleau C, Graham J & Canepari, M (2018). A novel multisite confocal system for rapid Ca2+ imaging from submicron structures in brain slices. J Biophotonics 11(3).

Fleidervish IA, Lasser-Ross N, Gutnick MJ & Ross WN (2010). Na+ imaging reveals little difference in action potential-evoked Na+ influx between axon and soma. Nat Neurosci 13, 852–860.

Goldwyn JH, Remme MWH & Rinzel J (2019). Soma-axon coupling configurations that enhance neuronal coincidence detection. PLoS Comput Biol 15, e1006476.

Hallermann S, de Kock CP, Stuart GJ & Kole MH (2012). State and location dependence of action potential metabolic cost in cortical pyramidal neurons. Nat Neurosci 15, 1007–1014.

Hu W, Tian C, Li T, Yang M, Hou H & Shu Y (2009). Distinct contributions of Na(v)1.6 and Na(v)1.2 in action potential initiation and backpropagation. Nat. Neurosci 12, 996–1002.

Jaafari N & Canepari M (2016). Functional coupling of diverse voltage-gated Ca(2+) channels underlies high fidelity of fast dendritic Ca(2+) signals during burst firing. J Physiol 594, 967–983.

Jaafari N, De Waard M & Canepari M (2014). Imaging Fast Calcium Currents beyond the Limitations of Electrode Techniques. Biophys J 107, 1280–1288.

Jaafari N, Marret E & Canepari, M (2015). Using simultaneous voltage and calcium imaging to study fast Ca2+ channels. Neurophotonics 2, 021010.

Katz E, Stoler O, Scheller A, Khrapunsky Y, Goebbels S, Kirchhoff F, Gutnick MJ, Wolf F & Fleidervish IA (2018). Role of sodium channel subtype in action potential generation by neocortical pyramidal neurons. Proc Natl Acad Sci USA 115, 7184–7192.

Kole MH, Ilschner S, Kampa B, Williams SR, Ruben PC & Stuart GJ (2008). Action potential generation requires a high sodium channel density in the axon initial segment. Nat Neurosci 11, 178–186.

Kole MH & Stuart GJ (2012). Signal processing in the axon initial segment. Neuron 73, 235–247.

Kushmerick MJ, Larson RE & Davies RE (1969) The chemical energetics of muscle contraction. I. Activation heat, heat of shortening and ATP utilization for activation-relaxation processes. Proc R Soc Lond B Biol Sci 174, 293–313.

Lamy CM & Chatton JY (2011). Optical probing of sodium dynamics in neurons and astrocytes. Neuroimage 58, 572–578.

Martina M & Jonas P (1997). Functional differences in Na+ channel gating between fast-spiking interneurones and principal neurones of rat hippocampus. J Physiol 505, 593–603.

Miyazaki K, Lisman JE & Ross WN (2019). Improvements in Simultaneous Sodium and Calcium Imaging. Front Cell Neurosci 12, 514.

Miyazaki K & Ross WN (2015). Simultaneous Sodium and Calcium Imaging from Dendrites and Axons. eNeuro. 2, pii: ENEURO.0092-15.2015.

Minta A & Tsien RY (1989). Fluorescent indicators for cytosolic sodium. J Biol Chem 264, 19449–19457.

Popovic MA, Foust AJ, McCormick DA & Zecevic D (2011). The spatio-temporal characteristics of action potential initiation in layer 5 pyramidal neurons: a voltage imaging study. J Physiol 589, 4167–4187.

Popovic M, Vogt K, Holthoff K, Konnerth A, Salzberg BM, Grinvald A, Antic SD, Canepari M & Zecevic, D (2015). Imaging Submillisecond Membrane Potential Changes from Individual Regions of Single Axons, Dendrites and Spines. Adv Exp Med Biol 859, 57–101.

Rasband MN (2010). The axon initial segment and the maintenance of neuronal polarity. Nat Rev Neurosci 11, 552–562.

Reynolds C, King MD & Gorman KM (2020). The phenotypic spectrum of SCN2A-related epilepsy. Eur J Paediatr Neurol 24, 117–122.

Roder P & Hille C (2014). ANG-2 for quantitative Na+ determination in living cells by time-resolved fluorescence microscopy. Photochem Photobiol Sci 13, 1699–1710.

Rush AM, Dib-Hajj SD & Waxman SG (2005). Electrophysiological properties of two axonal sodium channels, Nav1.2 and Nav1.6, expressed in mouse spinal sensory neurones. J Physiol 564, 803–815.

Scott RS, Henneberger C, Padmashri R, Anders S, Jensen TP & Rusakov DA (2014). Neuronal adaptation involves rapid expansion of the action potential initiation site. Nat Commun 5, 3817.

Spratt PWE, Ben-Shalom R, Keeshen CM, Burke KJ Jr, Clarkson RL, Sanders SJ & Bender KJ (2019). The Autism-Associated Gene Scn2a Contributes to Dendritic Excitability and Synaptic Function in the Prefrontal Cortex. Neuron 103, 673–685.

Stuart GJ & Sakmann B (1994). Active propagation of somatic action potentials into neocortical pyramidal cell dendrites. Nature 367, 69–72.

Vogt KE, Gerharz S, Graham J & Canepari, M (2011a). High-resolution simultaneous voltage and Ca^2+^ imaging. J Physiol 589, 489–494.

Vogt KE, Gerharz S, Graham J & Canepari, M (2011b). Combining membrane potential imaging with L-glutamate or GABA photorelease. PLoS ONE 6, e24911.

Wagnon JL & Meisler MH (2015). Recurrent and Non-Recurrent Mutations of SCN8A in Epileptic Encephalopathy. Front Neurol 6, 104.

Wimmer VC, Reid CA, So EY, Berkovic SF & Petrou S (2010). Axon initial segment dysfunction in epilepsy. J Physiol 588, 1829–1840.

Xiao Y, Jackson JO 2^nd^, Liang S & Cummins TR (2011). Common molecular determinants of tarantula huwentoxin-IV inhibition of Na+ channel voltage sensors in domains II and IV. J Biol Chem 286, 27301–27310.

Yin L, Rasch MJ, He Q, Wu S, Dou F & Shu Y (2017). Selective Modulation of Axonal Sodium Channel Subtypes by 5-HT1A Receptor in Cortical Pyramidal Neuron. Cereb Cortex 27, 509–521.

Yue C, Remy S, Su H, Beck H & Yaari Y (2005). Proximal persistent Na+ channels drive spike afterdepolarizations and associated bursting in adult CA1 pyramidal cells. J Neurosci 25, 9704–9720.

Zylbertal A, Kahan A, Ben-Shaul Y, Yarom Y & Wagner S (2015). Prolonged Intracellular Na+ Dynamics Govern Electrical Activity in Accessory Olfactory Bulb Mitral Cells. PLoS Biol 13, e1002319.

